# Building Bridges Belowground: *Pythium oligandrum* En-hances Rhizosphere Network Connectivity

**DOI:** 10.64898/2026.10.22.761010

**Authors:** Natalia Ramírez, Christian B. Andersen, Maja Brus-Szkalej, Linnea J. Stridh, Åsa Lankinen, Laura J. Grenville-Briggs

## Abstract

Biological control agents (BCAs) offer a potential sustainable alternative to fungicides for crop disease management, but their effects on resident plant microbiomes and microbial interactions remain poorly understood. Here, we investigated the effects of the BCA *Pythium oligandrum* on disease control and the rhizosphere microbiome of starch potato cv. Kuras in a field trial. We compared a laboratory strain and a commercial formulation (Polygandron) with commonly used fungicides during early blight development. The laboratory strain suppressed early blight at plant senescence, but this effect did not persist until harvest, whereas fungicide treatment provided near-complete disease suppression throughout the growing season. *P. oligandrum* caused transient changes in rhizosphere fungal community composition at flowering, while bacterial community richness varied among sampling times in untreated controls. However, bacterial co-occurrence network analysis revealed pronounced treatment-specific changes in microbial connectivity. Application of the laboratory strain increased network connectivity, resulting in almost a threefold increase in mean degree and greater interconnectedness among bacterial taxa. In contrast, Polygandron and fungicide treatments reduced mean degree, indicating reduced connectivity within the bacterial network. Fungicide treatment additionally altered network organisation, including increased betweenness centrality and changes in modularity. Together, our results show that *P. oligandrum* can modify microbial interactions in the potato rhizosphere without substantially altering overall bacterial community composition, highlighting the value of integrating microbial network structure with community-level analyses when evaluating the ecological effects of BCAs.

**Lay summary:** Biological control agents offer a more sustainable alternative to fungicides for managing crop diseases, but their effects on beneficial and resident microbes are not fully understood. We investigated how *Pythium oligandrum*, a potential biological control agent, affected early blight and the microbial communities around the roots of starch potato plants grown in a field trial. We compared a laboratory strain and a commercial formulation (Polygandron) with conventional fungicide treatment.

The laboratory strain reduced early blight symptoms towards the end of the growing season, but this effect did not persist until harvest. Fungicides provided more consistent disease control throughout the season. While treatments had limited effects on the overall composition of bacterial communities, they changed the patterns of associations between bacterial groups. The laboratory strain almost tripled the average number of associations per bacterial group, whereas Polygandron and fungicide treatments reduced this connectivity. Fungicides also changed how these associations were organised.

These findings show that biological control agents can influence relationships among root-associated microbes even when the overall bacterial community changes little. Understanding these microbial networks alongside community composition could help improve the evaluation of biological control strategies and support the development of more sustainable crop protection practices.

**Teaser text:** Biological control with *Pythium oligandrum* reshapes bacterial network connectivity in the potato rhizosphere, revealing microbial changes that remain hidden when community composition is considered alone.

## Introduction

Potatoes (*Solanum tuberosum* L.), ranked fourth among the largest food crops globally after rice, wheat, and maize [1], are a nutritious source of high-quality protein, energy, and essential minerals and vitamins, as well as an important source of starch. However, potato growth can be negatively affected by soil-borne pathogens, leading to significant yield and quality losses [2]. An example of a serious soil-borne pathogen is *Alternaria solani*, the causal agent of the disease early blight, which in severe cases, causes leaves to dry and shed rapidly, ultimately leading to the wilting and death of the entire plant [3]. Early blight is an increasing global problem that can cause yield losses of up to 30% in potato crops and affect the quality of the tubers, in particular in potato cultivars used for starch production [4,5]. Fungicides are traditionally used to control potato early blight [6]. However, fungicide resistance is emerging rapidly in *A. solani* populations [7,8], leading to vulnerable crop production and necessitating the search for alternative disease control methods [6].

The EU has implemented directives 2009/128 and 2019/782 to reduce the impact of large-scale agriculture, emphasizing integrated pest management (IPM) as a sustainable alternative to synthetic chemical controls [9]. A key IPM strategy is the use of biocontrol agents (BCAs), particularly microbial BCAs, which can suppress plant diseases while maintaining soil fertility and beneficial microbiomes [10,11]. Given the growing recognition of the plant microbiome’s role in crop health [12–15], it is crucial to assess how BCAs influence microbiome composition during disease control [16]. However, most studies on the potato rhizosphere have focused on temporal dynamics, genotypes, cropping systems, and plant development stages [17–20].

The BCA *Pythium oligandrum* has documented antagonist and mycoparasitic abilities against a wide range of hosts including plant pathogenic *Pythium* spp.[21], *Phytophthora* spp. [22–24], ascomycetes and basidiomycetes [25]. *P. oligandrum* exhibits key biocontrol traits, including rapid growth and induction of pathogen resistance in host plants [26,27]. Recently, we demonstrated that *P. oligandrum* reduced early blight in greenhouse-grown cv. Desiree potatoes and, in combination with Serenade®, decreased disease pressure in field-grown cv. Kuras potatoes [28]. However, its impact on the rhizosphere microbiome in the presence of disease remains unknown.

In this study, we investigated the impact of *P. oligandrum* on the rhizosphere microbiome of field-grown potato plants challenged with *A. solani*. For clarity, throughout this paper, *P. oligandrum* refers to the laboratory strain, whereas Polygandron refers to the commercial formulation of *P. oligandrum.* Given its broad host range and mycoparasitic activity against multiple fungal hosts [26,59,60], we hypothesized that *P. oligandrum* application would reduce fungal ASV richness and alter rhizosphere microbiome diversity and structure. We compared a laboratory strain and the commercial formulation Polygandron WP before and after *A. solani* infection, alongside a fungicide treatment (Narita + Propulse) as a positive disease-control treatment. Microbial co-occurrence networks provide insights into potential microbial associations and community organisation, complementing measures of diversity and composition. Species co-occurrence patterns are generally shaped by three main ecological processes: biotic interactions, environmental filtering, and dispersal limitation. [29,30]. However, how BCAs influence rhizosphere microbial networks, particularly under field conditions in potato, remains poorly understood. Disease suppression and rhizosphere microbiome responses were assessed using amplicon sequencing, including community composition, richness, core microbiome structure, and microbial co-occurrence networks.

We hypothesized the following:

i) Treatment with the BCA *P. oligandrum* against *A. solani* decreases early blight disease in starch potatoes.
ii) The effects of *P. oligandrum*, Polygandron and fungicide treatments on rhizosphere microbiome composition, diversity and network interconnectivity change over the course of the growing season.
iii) Treatment with *P. oligandrum* and synthetic fungicides influences rhizosphere microbiome composition, diversity and network interconnectivity in starch potatoes infected with *A. solani*, with different effects depending on the treatment applied.

## Materials and methods

### Study site and experimental treatments

The field trial was conducted in Sweden at the Swedish University of Agricultural Sciences (SLU) campus experimental garden, in Alnarp (55°66′106.2″N 13°08′198.9″E) in 2021. No potato crops had been cultivated in the area prior to the experiment, and rarely in the surroundings, reducing the probability of contamination.

Starch potato plants were artificially inoculated with *A. solani*. The experimental layout was done in a randomized block design, with 3 blocks and 4 treatments in each block with a random selection of the plants for microbiome analyses. The field trial was managed and fertilized following standards set by the Swedish Rural Economy and Agricultural Societies, with minor modifications. The fertilizers used were kindly supplied by YARA®, and were applied before planting, additional manganese fertilizers were applied throughout the growing season. For an overview of the management, agronomic inputs, treatments and data collection please see Table 1.

**Table 1.** Overview of field trial management, fertilization, early blight treatments, sampling, and disease assessment throughout the 2021 growing season. The table summarises the timing, treatment groups, and procedures applied across the experimental plots, including rhizosphere and bulk soil sampling. Rhizosphere and bulk soil samples were collected before inoculation with *Alternaria solani*.

| Activity | Treatment / group | Timing | Details |
| --- | --- | --- | --- |
| Trial management | All plots | 20 May 2021 | Seed tuber planting |
|  | All plots | 27 May 2021 | Plant emergence |
|  | All plots | Throughout trial | Watering when needed |
|  | All plots | 27 May; 8, 22 Jul; 5, 20 Aug; 2 Sep 2021 | Manual weeding |
| Fertilization | All plots | Before planting | YaraMila PROMAGNA (11-5-18), 1230 kg ha<sup>-1</sup> |
|  | All plots | 1, 8, 22 Jul; 20 Aug 2021 | YaraVita MANTRAC PRO, 200 L ha<sup>-1</sup> |
| Early blight treatment | Fungicide | 8, 22 Jul; 5, 20 Aug; 2 Sep 2021 | Narita (0.3 L ha<sup>-1</sup>) + Propulse (0.40 L ha<sup>-1</sup>); spray volume 300 L ha<sup>-1</sup> |
|  | Polygandron WP® | 8, 22 Jul; 5, 20 Aug; 2 Sep 2021 | 5 × 10<sup>5</sup> oospores g<sup>-1</sup>; 300 L ha<sup>-1</sup> |
|  | P. oligandrum CBS strain | 8, 22 Jul; 5, 20 Aug; 2 Sep 2021 | 2.5 × 10<sup>4</sup> oospores mL<sup>-1</sup>; 300 L ha<sup>-1</sup> |
| Rhizosphere sampling | Untreated control | 1 Jul*; 8 Jul; 5 Aug; 2 Sep 2021 | No early blight treatment |
|  | P. oligandrum-treated | 1 Jul*; 8 Jul; 5 Aug; 2 Sep 2021 | Rhizosphere samples |
|  | Polygandron-treated | 1 Jul*; 8 Jul; 2 Sep 2021 | Rhizosphere samples |
|  | Fungicide-treated | 1 Jul*; 8 Jul; 2 Sep 2021 | Rhizosphere samples |
|  | Untreated control | 1 Jul*; 5 Aug; 2 Sep 2021 | Rhizosphere samples |
| Early inoculation and blight scoring | Bulk soil | 1 Jul*; 2 Sep 2021 | Bulk soil samples |
|  | All treatment groups | 1 Jul 2021 | Inoculation with A. solani |
|  | All treatment groups | 22, 29 Jul; 5, 20, 27 Aug; 2, 9, 17, 24 Sep; 1 Oct | Disease scoring; first symptoms recorded 22 Jul |
|  | All treatment groups | 17 Sep; 24 Sep; 1 Oct 2021 | Disease scoring included defoliation |
| Plant height | All treatment groups | 1, 8, 15, 22, 29 Jul; 5, 13, 20, 27 Aug; 2, 9, 17, 24 Sep; 1 Oct | Measured at each scheduled assessment date |

\* Rhizosphere and bulk-soil samples were collected before inoculation with A. solani.

The seed tubers of potato starch cv. Kuras were kindly provided by Lyckeby SSF®. This specific cultivar was chosen based on its high agronomic performance and wide use in the Scandinavian region. The potatoes were planted with a row distance of 0.75 m and a planting distance of 0.38 m.

### Pathogen inoculation and disease assesment

*A. solani* (AS112), originally isolated from a field in southern Sweden [31] was cultivated on V8-medium at 22°C and 12h near visible ultraviolet light for 14 days. Afterwards, half of a 9 cm petri dish was kneaded into 150 g of double autoclaved barley grain in a tray soaked with 80 ml of sterile distilled water. The bag was incubated at 22°C and 12h near-visible ultraviolet light for two weeks, then turned over and kneaded again and finally incubated for additional 4 weeks at the same conditions. Five grams of *A. solani* inoculated barley grains/m^2^ were evenly scattered between the potato rows.

Early blight scoring was done weekly, starting one week post inoculation up until harvest following Duarte *et al.* [32]. Early blight severity was assessed as the leaf area covered by *A. solani* lesions. Treatment effects were evaluated using the area under the disease progress curve (AUDPC), calculated according to Shaner [33].

### Treatments, inoculum production and amendment

The four treatments were *Pythium oligandrum* fresh oospore preparation (CBS-strain 530.74), *P. oligandrum* commercial preparation as (Polygandron WP ®), standard commercially recommended fungicides (Narita® and Propulse®) and untreated, but *A. solani* inoculated controls. The inoculum preparation and application rate were identical to those described in Andersen *et al.* [34], with a final concentration of 2.5 × 10 oospores mL ¹ applied at 300 L ha ¹. Polygandron WP, containing 5×10^5^ oospores per gram, was diluted in water at the trial site to create a solution of 200 g/ha. The solution was applied at a rate of 300 L per hectare as recommended by the manufacturer. As a positive control we used a fungicide combination applied at the same time points as the BCAs and in the same dosage avoiding any spill over to adjacent plants (Table 1).

### Nondestructive rhizosphere sampling

Rhizosphere samples were collected using a soil core driller within 5 cm of each plant, following Andersen *et al*.[34]. Rhizosphere sampling was done four times during the cultivation period: before inoculation of *A. solani* (1st of July 2021), after around 6 hours of treatment inoculation (8th of July 2021), at the flowering stage (5th of August 2021) and, at senescence (2nd of September 2021). Three plants per block per treatment were sampled. For examination of the microbiome, we prioritized *P. oligandrum* and omitted Polygandron– and fungicide-treated plants at the flowering stage, since we wanted to focus our efforts on the interaction of the live biocontrol agent with the native microbial community. Please see Table 1 for the sampling design.

### Rhizosphere DNA extraction and sequencing

The frozen soil cores were thawed at ambient temperature and biological replicates were pooled together. A 0.25 g portion of each soil core was used for DNA extraction with the DNeasy PowerSoil Pro Kit (QIAGEN), following Andersen et [34]. The sequencing was performed by Illumina MiSeq V3 at LGC Genomics® in Germany obtaining approximately 60 KB reads. The primers sets used were 799F-1115R for bacteria and FITS7-ITS4 for fungi. Two different predefined mock communities from ZymoBIOMICS:Microbial Community Standard Biosite-D6310 and BioSite-D6306 were used as reference for sequencing perfor-mance and potential cross contamination exclusion. Additionally, to detect unintended contamination from the DNA extraction procedure we extracted blank controls from the Qiagen DNeasy PowerSoil Pro kit. The raw sequencing reads and associated metadata have been deposited in Zenodo and are available at https://doi.org/10.5281/zenodo.23471271.

### Statistical analysis

Disease quantification was analysed using RStudio (version 4.1.2 2022), using rstatix [35], and Car [36] packages. Boxplots were generated with ggplot2 [37]. Treatment effectiveness was assessed via analysis of variance (ANOVA) with square root transformed AUDPC as the response variable and treatment and block as predictors.

To evaluate the impact of treatments on microbiome alpha-diversity, we conducted ANOVA and Tukey HSD test. Differences in microbiome community composition (beta diversity) were performed using PERMANOVA with 999 permutations [38]. Non-metric multidimensional scaling (NMDS) based on the Bray-Curtis dissimilarity matrix was performed to visualize differences between *P. oligandrum* fresh oospore treatment and control plants. All microbiome data were analysed and visualized in RStudio, using a combination of phyloseq package [39] and Microeco package [40].

To investigate potential differential treatment effects on the fungal to bacterial abundance, the Fungal:Bacterial ratio was calculated for each sample using the R environment (v.4.x) and the phyloseq package [39]. The Fungal:Bacterial ratio was then derived by dividing the total fungal abundance by the total bacterial abundance for each sample and added to the sample metadata for downstream visualization and statistical analysis. For statistical analysis, differences in Fungal:Bacterial ratios among treatments were assessed using one-way ANOVA, followed by Tukey’s HSD post hoc test for pairwise comparisons.

### Co-occurrence networks construction

Microbial co-occurrence networks were constructed separately for each treatment and sampling date to evaluate treatment-specific microbial interactions. ASVs present in fewer than 90% of samples within each group were removed prior to analysis to reduce the influence of rare taxa. Raw ASV count data from phyloseq objects [39] were used after pruning zero-sum taxa and samples. Pairwise associations between ASVs were calculated using Spearman’s rank correlation coefficient with the rcorrfunction from the Hmisc package [41]. Only strong and significant correlations (|ρ| ≥ 0.6, p < 0.01) were retained and used as network edges representing positive and negative microbial associations.

Network topological properties, including degree centrality, modularity, betweenness centrality, clustering coefficient, and average degree, were calculated using the igraph package [42]. Modularity, betweenness and clustering coefficient were calculated from positive-only subnetworks. Because fungal co-occurrence networks contained substantially fewer nodes and edges than their bacterial counterparts, degree and betweenness analyses were not conducted for the fungal community. Statistical analyses were restricted to within-treatment comparisons due to substantial variation in the initial soil microbiome arising from natural heterogeneity.

### Relative abundance and community composition analysis

Relative abundance analyses were performed in R using the phyloseq and tidyverse ecosystems. Raw ASV (Amplicon Sequence Variant) counts were normalized using Total Sum Scaling (TSS), where counts in each sample were divided by the total number of reads in that sample using transform_sample_counts() in phyloseq. For visualization, ASVs were taxonomically aggregated at the phylum or genus level using tax_glom(). To reduce visual complexity, only the 10-15 most abundant taxa were displayed individually, while remaining taxa were grouped as “Other.”

## Results

### *P. oligandrum* treatment against *A. solani* reduces potato early blight during senescence

To investigate whether *P. oligandrum* and synthetic fungicides control early blight, we characterized the efficacy of the biocontrol agent (lab strain and the commercial product Polygandron) along with synthetic (Narita and Propulse) in potato (cv. Kuras). Disease progression was compared between untreated control, the two *P. oligandrum* treatments, and the fungicide treatment at the senescence and at tuber harvest. *P. oligandrum* treatment resulted in a significant decrease of the disease at the senescence stage of the potato plants (p = 0.03), however, this effect did not persist when the tubers were harvested one month later (Fig. 1). Interestingly, the commercial formulation of *P. oligandrum*, showed a tendency towards disease suppression, but was not significantly different from the control. On the contrary, the fungicide treatment almost completely inhibited the early blight disease at senescence (p < 0.001) (Fig. 1A) and the effect persisted up until the harvest of the potato tubers (p < 0.001) (Fig. 1B).

**Figure 1.**
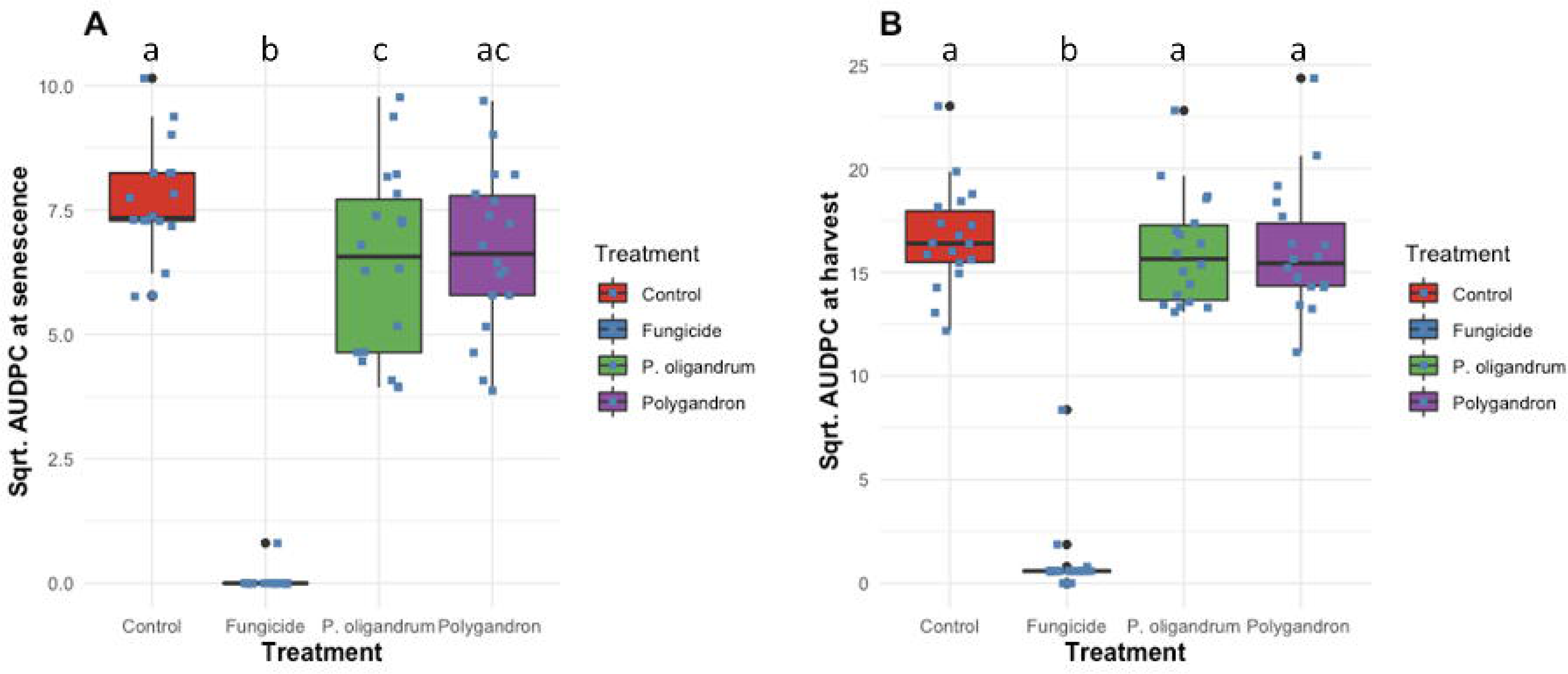
Square root transformed AUDPC (area under the disease progression curve) at (A), senescence (2nd of September) and (B), harvest (1^st^ of October). Untreated control plants infected with early blight are compared to treatments with fungicide, *P. oligandrum* or Polygandron, a commercially formulated oospore powder from *P. oligandrum*. Different lowercase letters indicate significant (p < 0.05) differences

### *P. oligandrum* treatment for early blight temporarily alters the rhizosphere microbiome

We sequenced the rhizosphere microbiome for all treatments at three time points: before *A. solani* inoculation (1st of July), post-inoculation (8^th^ of July), and at senescence (2nd of September) to assess treatment effects over the growing season. Additionally, we sequenced *P. oligandrum* and control treatments also at flowering (5^th^ of August) (Table 1). In our first analysis, we considered whether the early blight suppression observed with the *P. oligandrum* laboratory strain at the senescence (Fig. 1A) was associated with changes in the rhizosphere microbiome compared to control treatment (Table 1). Bacterial ASV abundance did not differ significantly between treatments (Fig. 2A-D), whereas fungal ASV abundance was significantly higher in *P. oligandrum*-treated plants at flowering (Fig. 2E-H). Considering all investigated treatments and time points, bacterial beta diversity differed significantly between control and both Polygandron (adjusted *p* = 0.009) and fungicide treatments (adjusted *p* = 0.006; Fig. 3C), while fungal beta diversity differed significantly between control and *P. oligandrum*-treated plants at flowering (*p* = 0.022; Fig. 3F). However, the Fungal:Bacterial ratio did not change when comparing the early and senescence sampling periods. (S.F.1).

**Figure 2.**
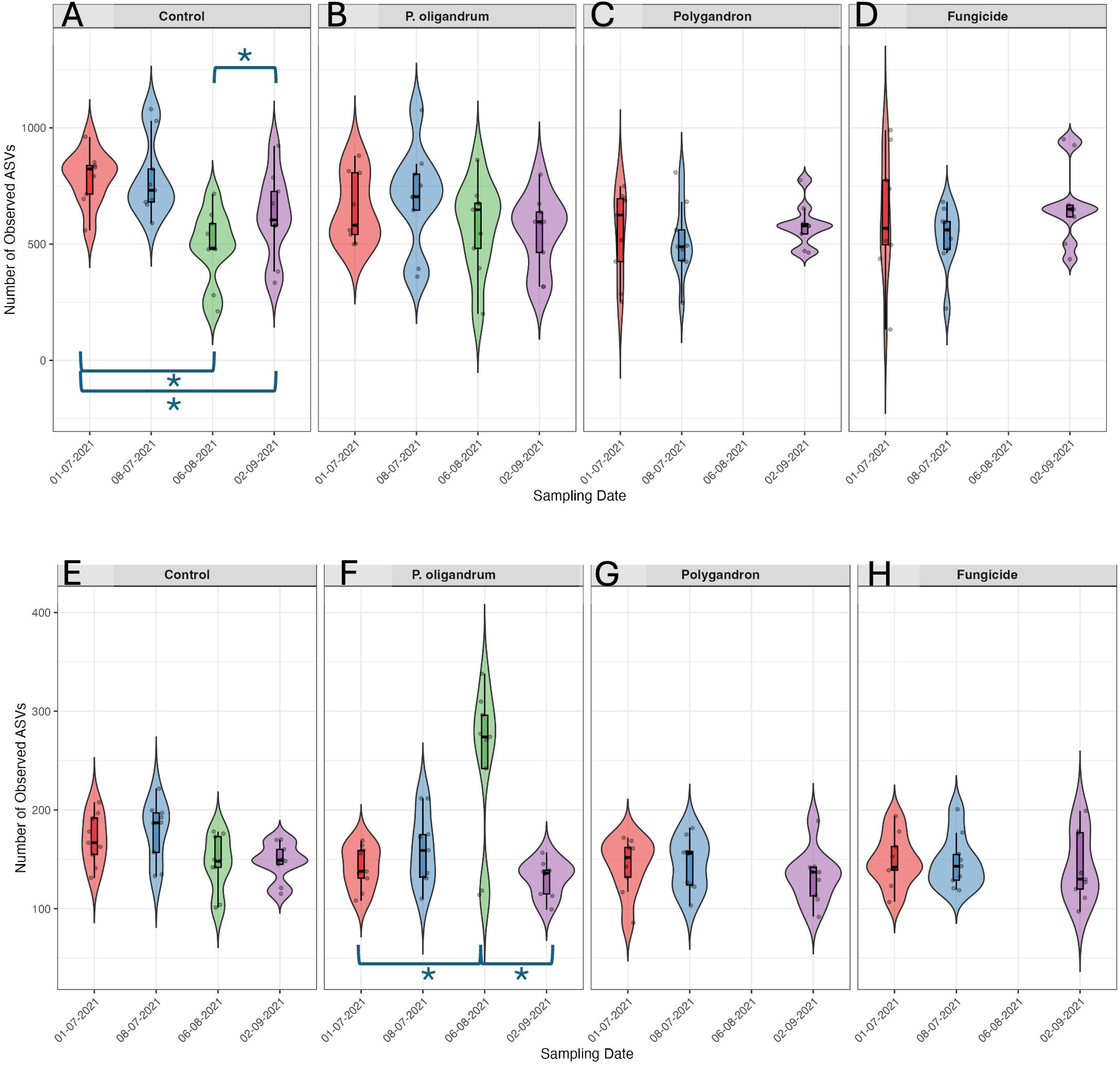
Observed ASV (Amplicon Sequence Variant) A-D bacterial ASVs; E-H fungal ASVs. Asterisks show statistical differences among sampling times (p < 0.

**Figure 3.**
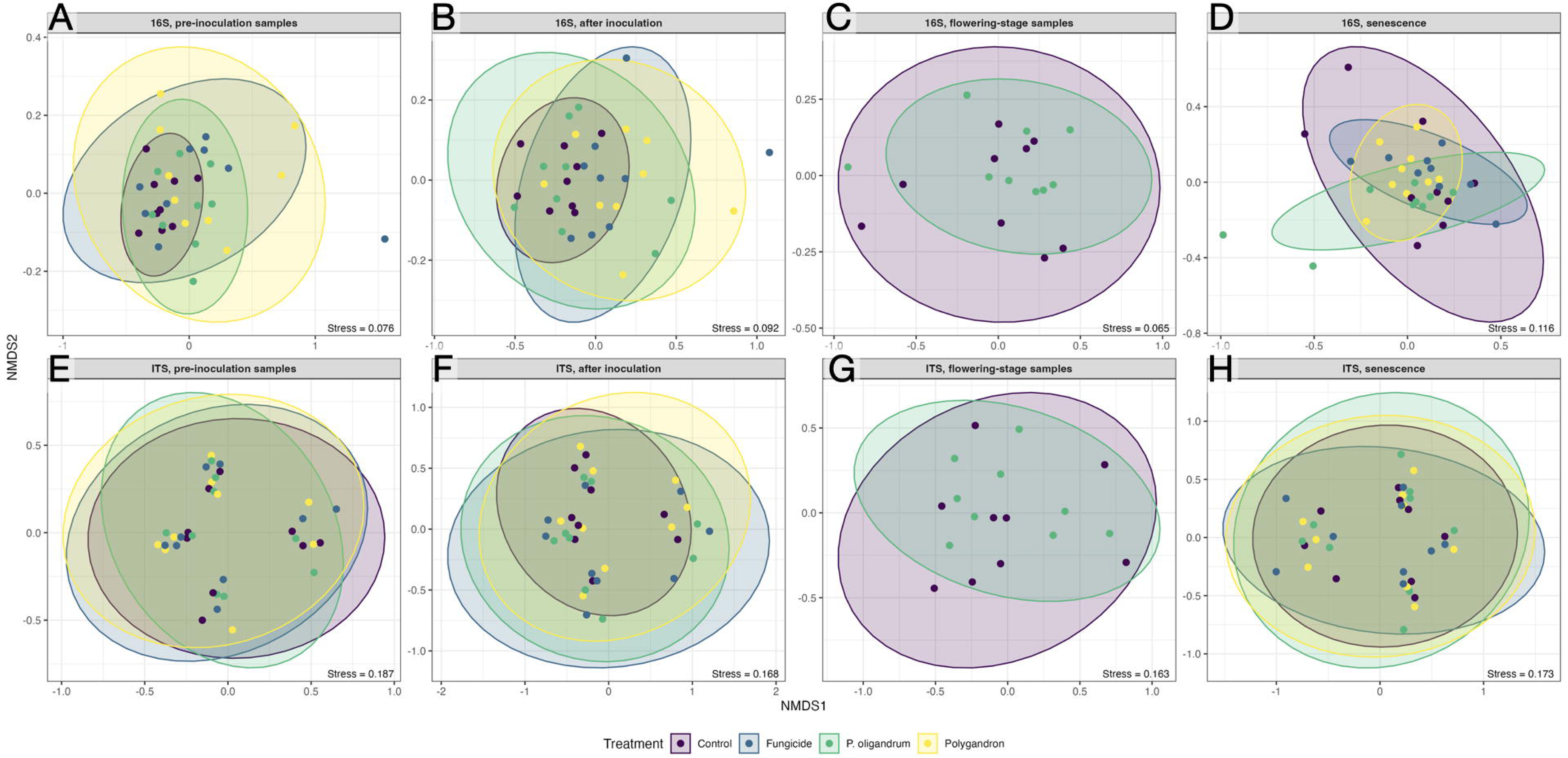
Non-metric multidimensional scaling (NMDS) plots illustrating beta diversity of bacterial (a–d) and fungal (e–h) communities in potato rhizosphere samples under *A. solani* infection, comparing untreated control plants with plants treated with the commercial formulation Polygandron WP® or the laboratory strain of *P. oligandrum*. Panels (a, e) show pre-inoculation samples (1^st^ of July), (b, f) flowering-stage samples (5^th^ of August), (c, g) samples collected one week after inoculation (8^th^ of July), and (d, h) samples collected at senescence (2^nd^ of September). Significant differences between treatments were assessed using PERMANOVA and are indicated by an asterisk (*p* < 0.05).

### Differences in the rhizosphere microbiome over the growing season

In an analysis of all treatments across the investigated time points, only the untreated control and *P. oligandrum* treatments differed significantly in fungal diversity at flowering, and the bacterial beta diversity between the control and Polygandron and fungicide treatments (Fig. 3). The fungal:bacteria ratio was higher in *P. oligandrum*treated plants than in controls at flowering (p < 0.01) and after inoculation (p < 0.05; S.F.1). Bacterial ASVs were largely shared across treatments and time points, indicating a predominance of generalists (S.F.2 A-D and S.F.3 A-D), whereas fungi showed more unique ASVs (S.F.2 E-H and S.F.3 E-H). Abundance-occupancy analysis confirmed a strong dominance of bacterial generalists, with only three specialists detected under *P. oligandrum*. Fungal communities contained more specialists than generalists. The bacterial abundance– occupancy relationship was steeper in the control than in the fungicide and Polygandron treatments, indicating a stronger association between ASV abundance and occupancy in untreated samples. The *P. oligandrum* treatment also showed a steeper slope than the fungicide treatment, whereas no significant differences were detected between the control and *P. oligandrum* or among the remaining treatment pairs. No significant differences among treatments were observed in the fungal microbiome (S.F.2; S.F.3).

Differential-abundance analysis identified some bacterial taxa that differed between treatments at specific stages of the experiment. Before inoculation, *Luteimonas* was more abundant in the *P. oligandrum*, Polygandron and fungicide treatments than in the control, whereas *Paenarthrobacter* was less abundant in Polygandron-treated plants. After inoculation, *Nocardioides* was less abundant in fungicide-treated plants, and it was also less abundant in *P. oligandrum*-treated plants at flowering. At senescence, *Actinomadura* was more abundant, whereas *Streptomyces* was less abundant, in fungicide-treated plants compared with the control. These taxa have been associated with diverse ecological functions, including plant growth promotion, nutrient acquisition, pathogen antagonism and secondary-metabolite production [43]. However, the differences detected at pre-inoculation should be considered baseline variation rather than treatment effects (S.F.4).

Fungal differential-abundance analysis revealed more pronounced treatment– and stage-specific shifts than bacterial analysis. After inoculation, *P. oligandrum* and Polygandron were associated with increased abundances of taxa such as *Tetracladium*, *Hymenoscyphus* and *Cladosporium*, while Polygandron also reduced *Vishniacozyma victoriae* and *Lachnum pygmaeum*. At flowering, *P. oligandrum* produced the largest number of differences, including increases in *Cladosporium*, *Apiotrichum* and *Gibberella/Fusarium*, and decreases in *Debaryomyces* and *Plectosphaerella*; at senescence, treatment-associated changes involved taxa linked mainly to saprotrophic, symbiotic or plant-associated lifestyles, including *Mycosphaerella*, *Paxillus* and *Roussoella* (S.F.5). Several reduced taxa, including *V. victoriae*, *Candida sake* and *Debaryomyces hansenii*, have documented antagonistic activity against plant pathogens [44–46], whereas *Fusarium* taxa include organisms with pathogenic potential, including important pathogens of potato [47].

Using a non-destructive sampling approach, we assessed whether rhizosphere samples could be distinguished from bulk soil at two time points. Before *A. solani* inoculation, bacterial beta diversity and both alpha and beta fungal diversity differed significantly between rhizosphere and bulk soil. At senescence, fungal alpha diversity and bacterial beta diversity differed significantly, while the other measures did not (S.F.6 and S.F.7). Overall, non-destructive sampling captured a microbiome distinct from bulk soil in potato.

### *P. oligandrum* increases bacterial rhizosphere microbiome interconnection

Despite limited treatment effects on bacterial diversity, we investigated whether inoculation altered rhizosphere network connectivity. Bacterial co-occurrence networks were analysed using node degree, betweenness centrality and modularity. Node degree describes the number of connections associated with each microbial taxon, betweenness centrality indicates the extent to which a taxon connects different parts of the network, and modularity reflects the degree to which the community is organised into distinct groups of closely interacting taxa. After inoculation with *P. oligandrum*, the rhizosphere bacterial co-occurrence network showed a 181% increase in average degree, indicating a higher number of associations between bacterial taxa (Figure 4). In contrast, Polygandron reduced the average degree by approximately 38%, while fungicide treatment reduced it approximately 66.6%, suggesting a lower level of microbial connectivity. Fungicide treatment also increased betweenness centrality, a measure of the extent to which taxa act as connecting nodes between different parts of the network, suggesting a more dispersed network structure (Fig. 4A). These results are supported by the modularity that is reduced around 38% after *P. oligandrum* treatment while it is doubled after fungicide treatment. In contrast, cooccurrence networks in the untreated control remained relatively stable (around 14% increase) between *A. solani* inoculation and a week after with the other treatments were added, showing fewer changes than those in the other treatments (Figure 4A; S.T.1).

**Figure 4.**
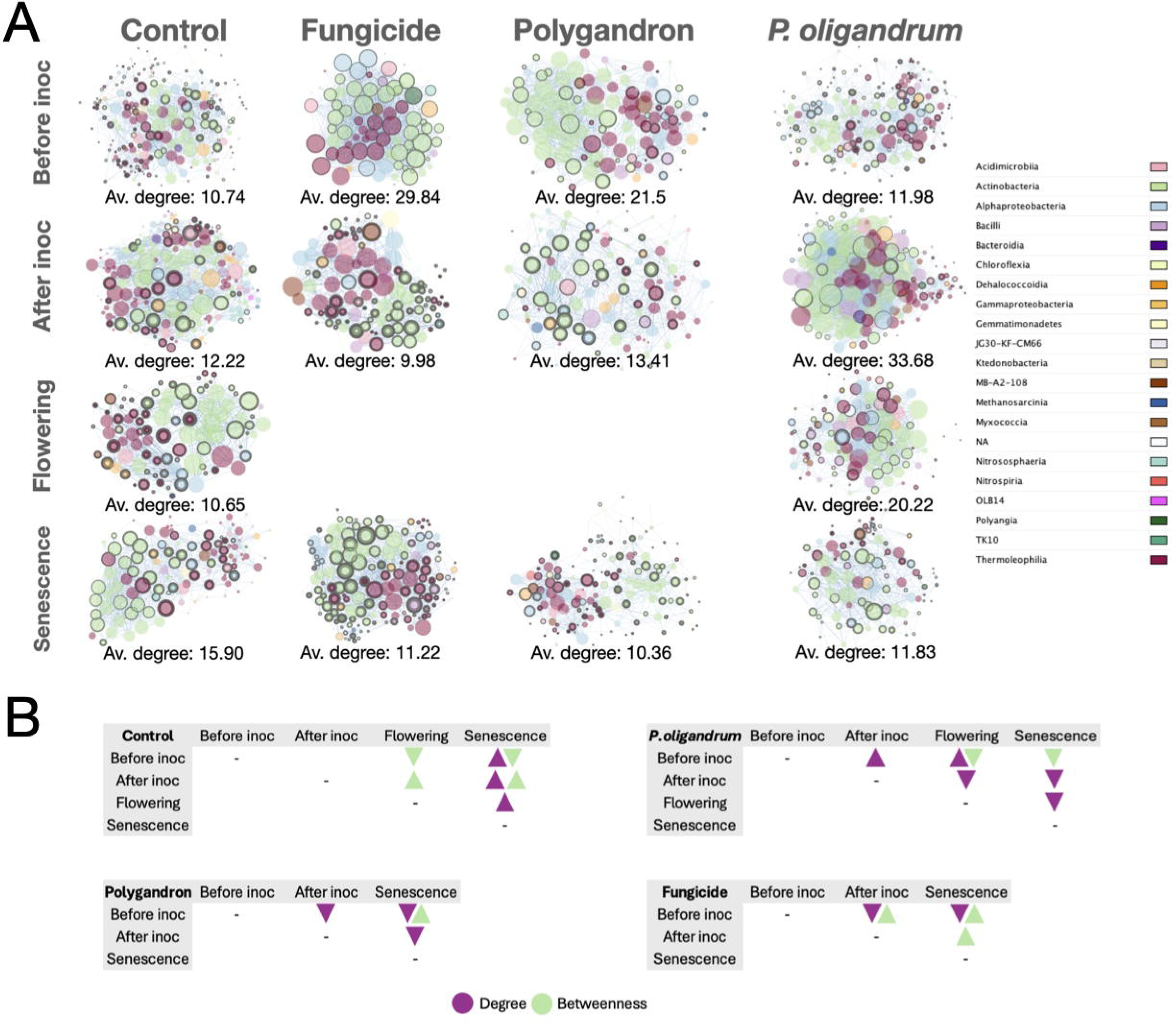
a) Co-occurrence network of the bacterial microbiome at different treatments (x axis) and sampling times (y axis). The color represent different taxonomical class while the size of the nodes (circles) represents the degree, and the border of the circles shows the betweenness. We additionally report the average degree (2edge/node) as a simple proxy of network connectivity.b) Pairwise comparisons of network topology metrics among sampling stages based on Dunn’s post hoc test. Significant differences in degree (purple) and betweenness centrality (green) are shown for each treatment. Upward and downward triangles indicate significantly higher and lower values, respectively, in the sampling stage shown in the column relative to the stage shown in the row.

The taxa exhibiting the highest degree centrality prior to treatment application were predominantly affiliated with Actinobacteriota and Alphaproteobacteria. Across all treatments, several taxa, including *Nocardioides*, *Solirubrobacter*, and *Cellulomonas*, consistently ranked among the most highly connected members of the network, indicating their potential role as core taxa within the soil microbiome (S.T.2).

In the control treatment, the composition of highly connected taxa remained relatively stable throughout the growing season. While *Nocardioides*, *Solirubrobacter*, and *Cellulomonas* consistently displayed high degree values, other taxa such as *Gaiella*, *Marmoricola*, *Longivirga*, and *Lapillicoccus* varied in their relative importance across sampling dates, reflecting natural seasonal succession within the microbial community (S.T.2).

The fungicide treatment produced a contrasting response. Following application, average degree decreased substantially, indicating a reduction in network interconnectivity. Taxa affiliated with *Thermoleophilia* became increasingly important, while members of Acidimicrobiia and Gemmatimonadota contributed more strongly to network structure. Although Rhizobiales remained present among the highly connected taxa, their relative importance decreased compared with other treatments. These shifts suggest that fungicide application selected for microbial communities dominated by oligotrophic and stress-tolerant taxa while maintaining a subset of rhizosphere-associated microorganisms. At senescence, the fungicide-treated community was characterized by highly connected taxa commonly associated with environmental stress tolerance, including *Blastococcus* [48] (S.T.2).

In contrast, inoculation with *P. oligandrum* induced a rapid restructuring of the bacterial network. Shortly after inoculation, network connectivity increased markedly, reaching the highest average degree observed among all treatments. Beneficial taxa such as *Bacillus* and *Tumebacillus* emerged as highly connected nodes, suggesting an important role in shaping community interactions following inoculation. During the flowering stage, network connectivity remained elevated; however, the degree distribution became less concentrated, with highly connected nodes distributed across a broader range of taxa. By plant senescence, overall connectivity declined, indicating a progressive relaxation of the initial network restructuring induced by *P. oligandrum* (S.T.2).

The commercial *P. oligandrum* formulation, Polygandron, produced an intermediate response between the untreated control and the pure *P. oligandrum* inoculation. Following Polygandron application, *Bacillus* became the most highly connected taxon, while members of Rhizobiales, particularly Xanthobacteraceae, emerged among the dominant hubs. In addition, cellulose-degrading taxa such as *Cellulomonas* displayed increased connectivity [49]. Although Actinobacteriota remained the dominant phylum throughout the season, overall network connectivity decreased after inoculation, suggesting a reorganization of co-occurrence associations rather than a general increase in interconnectivity (S.T.2).

*Pseudonocardia* was detected only following Polygandron and *P. oligandrum* inoculation. This genus has been reported as a plant growth-promoting bacterium with both antifungal and antibacterial activities, suggesting a potential role in plant health and disease suppression [50]. Similarly, Ilumatobacteraceae and *Blastococcus* were detected exclusively after *P. oligandrum* inoculation. *Bacillus*, which is widely recognized for its biostimulant and biocontrol properties, was also detected following *P. oligandrum* inoculation. [46–48] (S.T.2). The enrichment of these taxa may indicate that *P. oligandrum* promotes the establishment of microbial communities associated with plant growth promotion and pathogen suppression. Notably, the taxa identified as key network hubs based on degree and betweenness centrality were not members of the *A. solani* or *P. oligandrum* taxa.

Analysis of network modularity revealed contrasting responses among treatments following inoculation. A pronounced decrease in modularity was observed after *P. oligandrum* inoculation, indicating a reduction in network compartmentalization and an increase in interactions among previously separated microbial groups. In contrast, fungicide application resulted in a substantial increase in modularity, suggesting the formation of more distinct microbial modules and a more compartmentalized network structure. The control treatment exhibited relatively stable modularity values throughout the growing season, reflecting natural temporal variation without major network reorganization. Similarly, the Polygandron treatment showed only minor changes in modularity over time, maintaining values comparable to those observed in the control (Figure 4.B). Because fungal cooccurrence networks contained substantially fewer nodes and edges than their bacterial counterparts, degree and betweenness analyses were not conducted for the fungal community.

## Discussion

In this study, we evaluated the efficacy of the biocontrol agent *P. oligandrum* and synthetic fungicides in controlling early blight in starch potatoes and assessed the impact of these treatments on the rhizosphere microbiome using metabarcoding. We found that *P. oligandrum* treatment significantly decreased early blight disease at the plant senescence but lost efficacy by harvest, while fungicides almost completely eradicated the disease. Limited changes were observed in rhizosphere alpha and beta diversity and the fungal-to-bacterial ratio. However, *P. oligandrum* induced a transient increase in bacterial network connectivity and reduced community compartmentalization, whereas fungicide treatment resulted in reduced connectivity and increased modularity, indicating contrasting effects on rhizosphere microbial network structure. In contrast rhizosphere bacterial interconnections were reduced by the application of either fungicides or Polygandron.

### Early blight disease control differs between *P. oligandrun* and synthetic fungicide treatments

*P. oligandrum* transiently suppressed early blight in starch potato, with an effect observed only at the senescence point, when disease typically begins to develop or remains at low levels. This suggests that *P. oligandrum* may be most effective under low disease pressure or during the early stages of disease development, as previously reported by Stridh *et al.* [28]. Similar effects have been reported for *B. amyloliquefaciens* [47] and for both *B. amyloliquefaciens* and *Trichoderma* spp. [54,55]. In the present study the commercially formulated Polygandron did not significantly reduce the disease at any time point. This may be related to the limited oospore germination that causes reduced colonization and proliferation in field conditions (personal communication, Thomas Rey, De Sangosse). In contrast, fungicide treatment achieved complete disease suppression at senescent stage, and this persisted up until harvest as previously shown in field conditions [28]. In conclusion, while our data suggest some early blight control by hand sprayed *P. oligandrum* in starch potato, its efficacy as a standalone is insufficient as a BCA. Future studies should optimize the application method, rate, timing, and formulation of *P. oligandrum* to enhance its efficacy as a BCA under field conditions.

### *P. oligandrum* treatment for early blight temporarily alters the rhizosphere microbiome composition and diversity

We also examined the effects of *P. oligandrum* and synthetic fungicides on the rhizosphere microbiome composition and diversity, as understanding plant-microbial interactions is essential for low-input management strategies, high-quality crops [56] and an increased yield [53–55]. Thus, unwanted changes in the potato microbiome due to BCA treatments are likely a disadvantage for overall plant health. At the flowering stage, *P. oligandrum* treatment did not significantly affect the alpha or beta diversity of the potato rhizosphere bacterial community. In contrast, the fungal community showed significant changes in ASV composition at flowering, distinguishing it from the same treatment at pre-inoculation and senescence. Given that *P. oligandrum* has a broad host range and preys on multiple hosts across [26,60,61], we predicted a reduction in the fungal ASVs after its application. While the beta diversity of fungal communities differed significantly between *P. oligandrum* treated and control plants at this stage, no significant effect was observed at the alpha-diversity level. The differential abundance analysis at the flowering stage, showed an impact on the fungal community by *P. oligandrum*, complementing our findings at the ASV level.

The effect of *P. oligandrum* on early blight at senescence could have influenced the rhizosphere microbiome, as early blight is known to alter bacterial and fungal community structure in potato [62]. However, we found no significant changes in alpha or beta diversity at this stage, consistent with some studies in tomato [63,64]. In contrast, transient changes in rhizosphere diversity have been reported following *B. amyloliquefaciens* and *Pseudomonas fluorescens* application [65,66], while other studies have found more persistent effects of *B. amyloliquefaciens* and *Trichoderma* spp.[63–68]. These contrasting findings suggest that microbiome responses depend on factors such as the BCA, host plant, soil, disease, dose, sampling strategy, and timing [73]. Although overall diversity remained largely unaffected in our study, changes in microbial associations may occur without detectable shifts in community diversity or composition. We therefore examined co-occurrence networks to investigate whether the different treatments altered patterns of potential microbial associations and community organisation.

### *Pythium oligandrum* promoted network connectivity, whereas fungicide application reduced network connectivity

*P. oligandrum* inoculation increased the average degree of the microbial co-occurrence network, whereas fungicide treatment and Polygandron inoculation reduced it. In contrast, the control treatment remained relatively stable across sampling dates. Similar effects of BCA application on microbial network topology have been reported by Cardoni et al. [74], who found that two biocontrol rhizobacteria differentially altered olive root-associated microbial networks despite limited changes in overall community composition. A higher average degree reflects a more interconnected microbial community, with taxa participating in a greater number of cooccurrence relationships. Such increased connectivity may contribute to greater network stability and functional redundancy within the microbial community [75,76]. Consistent with this potential link between network connectivity and soil health, Zheng et al. [77] associated a higher number of co-occurrence network edges with healthier roots and disease-suppressive soils than with diseased roots and disease-conducive soils. However, increased connectivity alone does not necessarily indicate greater stability or a healthier microbiome.

The lower average degree observed following Polygandron inoculation is consistent with its comparatively weaker beneficial effects relative to the laboratory strain of *P. oligandrum*. This interpretation is further supported by its limited impact on early blight suppression, as disease levels did not differ significantly from those observed in the control treatment. Together, these findings suggest that the reduced efficacy of Polygandron may be associated with a lower capacity to promote microbial network connectivity in the rhizosphere. This weaker effect could potentially be related to differences in formulation, inoculum characteristics, or other components of the commercial product. Further research is needed to elucidate the relationship between disease severity and microbial network connectivity.

In this study we used a combination of two fungicides: Narita containing difenoconazole which inhibits *A. solani* mycelia growth [78], reduces disease severity, increases tuber yield, and decreases soil bacterial diversity [79]; and Propulse containing Fluopyram, widely used for potato early blight, has been reported to affect both fungal and bacterial rhizosphere microbiomes. Previous studies have reported contrasting effects of fungicides on soil microbial communities. While Zhang et al. [80] found that fluopyram altered microbial community structure and function, Sun et al. [81] reported no negative effects on the microbiome. Similarly, in our study, fungicide treatment had no significant effect on rhizosphere microbiome alpha or beta diversity. However, following fungicide application, reduced average degree and increased betweenness indicated lower connectivity and greater reliance on bridging taxa. In contrast, increased average degree and reduced betweenness under *P. oligandrum* suggested a more densely interconnected network with connectivity distributed across more taxa.

The changes in network structure were accompanied by shifts in the identity and relative importance of highly connected bacterial taxa, providing potential insights into the mechanisms underlying these contrasting responses. Following *P. oligandrum* inoculation, *Bacillus* and *Tumebacillus* emerged as highly connected taxa. Members of these genera have been associated with plant growth promotion, biostimulation, and biocontrol activities [51,52], suggesting that their increased network importance may be relevant to the beneficial effects associated with *P. oligandrum*. Similarly, *Pseudonocardia* was detected following both *P. oligandrum* and Polygandron treatments and has previously been associated with plant growth-promoting and antimicrobial activities [50]. In contrast, the increasing importance of taxa associated with environmental stress tolerance following fungicide application may indicate stronger environmental filtering, favouring a subset of resilient taxa and contributing to the observed reduction in network connectivity. The intermediate response of Polygandron, including the increased importance of *Bacillus*, Xanthobacteraceae, and *Cellulomonas*, suggests that the commercial formulation may reproduce some, but not all, of the microbial restructuring associated with *P. oligandrum*. Overall, these findings indicate that treatment effects on the rhizosphere microbiome may involve changes not only in the abundance of individual taxa but also in their relative importance within the microbial network.

In conclusion, *P. oligandrum* reduced early blight symptoms in cv. Kuras at senescence, but suppression was weaker than with fungicides and did not persist until harvest. Thus, further optimization is needed to improve its efficacy as a biocontrol strategy. All treatments except the control highly altered bacterial rhizosphere network connectivity: *P. oligandrum* increased network degree and reduced modularity, whereas Polygandron and fungicide treatments reduced connectivity, with fungicide treatments also increasing betweenness and modularity. As *P. oligandrum* caused limited changes in rhizosphere diversity while increasing network connectivity, in our study, it may represent a relatively low-impact biological treatment. Further research should focus on optimizing BCA performance under field conditions as sustainable alternatives to synthetic fungicides.

## Author contributions CRediT statement

**Natalia Ramirez:** Methodology, Data Curation, Formal Analysis, Visualisation, Writing original draft, Writing review and editing; **Christian Andersen**: Conceptualisation, Investigation, Methodology, Formal Analysis, Visualisation, writing, original draft, writing review and editing; **Maja Brus-Skalej**: Investigation, Formal Analysis, Data Curation, writing review and editing; **Linnea Stridh**: Methodology, Investigation, writing review and editing; **Åsa Lankinen**: Conceptualisation, Funding acquisition, supervision, methodology, Formal Analysis, Writing review and editing; **Laura Grenville-Briggs**: Conceptualisation, Funding acquisition, Supervision, Methodology, Project Administration, Writing review and editing.

All authors read and approved the final version of the manuscript prior to submission.

## Funding

We gratefully acknowledge funding from: The Swedish Research council (grant nrs 2018-04354 and 2023-05529), The Carl Trygger foundation, The Swedish Research Council Formas (grant nrs 2019-00881 and 2024-02274), and SLU Grogrund. This project has also received funding from the European Union’s Horizon 2020 research and innovation programme, MSCA-ITN-2017 under grant agreement No 766048 to LGB, which supported CBA.

## Data availability

The raw sequencing reads and associated metadata have been deposited in Zenodo and are publicly available at https://doi.org/10.5281/zenodo.23471271.

## Supporting information

Visual abstract

## Acknowledgements

The authors would like to thank Adam Flöhr and Jan-Eric Englund, at Statistics @ SLU, who helped with statistics. We would like to thank Erland Liljeroth for his expertise in the pathosystem and on the contributions to study design. We further would like to thank Odlingsenheten @ SLU for the field management and fungicide application. In addition, we would like to thank Nicolas Delhomme and the SLUBI team @ SLU for bioinformatic support. We thank, YARA for supplying fertilizers for the field trial. We thank Lyckeby Starch SF for supplying the Tubers. We also thank Simon B. Lassen from the Department of Plant and Environmental Sciences, University of Copenhagen, for his assistance with the DADA2 analyses.

**S.F. 1. Fungi:bacteria ratio across treatments and sampling stages.** Violin plots with embedded boxplots show the distribution of fungi:bacteria ratios for bulk soil, untreated control, fungicide, *Pythium oligandrum*, and Polygandron treatments at four sampling stages: 1st of July (pre-inoculation), 8th of July (post-inoculation), 6th of August (flowering), and 2nd of September (senescence). Individual points represent biological replicates; boxplots indicate the median and interquartile range.

**S.F. 2** Abundance-occupancy analysis was performed in R using phyloseq and tidyverse. ASVs represented by more than one sequence were retained. Occupancy was calculated as the proportion of samples within each treatment in which an ASV was detected, while abundance was defined as its maximum relative abundance after Total Sum Scaling normalization.ASVs were classified as Generalists (present in ≥60% of samples), Specialists (maximum relative abundance >2% and present in <10 samples), or Neither. Abundance-occupancy plots were generated with occupancy on the x-axis and log10-transformed maximum relative abundance on the y-axis. To check that differences in sample size and sampling dates were not driving the results, we repeated the analysis using only the three dates sampled in every treatment. For each ASV, occupancy (the number of samples it was detected in out of the total) was modelled as a function of log10 mean relative abundance, treatment and their interaction. We used a binomial generalized linear mixed model (lme4) with random intercepts for ASV and for each observation. The interaction was tested with a likelihood-ratio test, and slopes were compared between treatments using Tukey-adjusted pairwise contrasts (emmeans). Panels a-d show bacterial (16S rRNA gene) communities: (a) Control, (b) *P. oligandrum*-treated, (c) Polygandron-treated, and (d) Fungicide-treated samples. Panels e-h show fungal (ITS) communities in the same treatment order: (e) Control, (f) *P. oligandrum*-treated, (g) Polygandron-treated, and (h) Fungicide-treated samples. Each point represents a single taxon, plotted according to its occupancy (x-axis; proportion of samples in which the taxon was detected) and maximum relative abundance (y-axis; log10 scale), corresponding to the highest relative abundance reached by that taxon in any individual sample within a group. Taxa were classified as generalists (blue) when present in ≥60% of samples within a group, indicating broad distribution across the community. Specialists (red) were defined as taxa reaching >2% relative abundance in at least one sample but detected in fewer than 10 samples within the group, indicating high abundance but restricted occurrence. Taxa not meeting either criterion were classified as neither (grey).

**S.F. 3.** UpSet plots were generated using UpSetR to identify ASVs unique to or shared among Control, *P. oligandrum*, Polygandron, and Fungicide treatments at each sampling date. Panels (a-d) show the 16S dataset at (a) pre-inoculation, (b) after inoculation, (c) flowering, and (d) senescence. Panels (e-h) show the ITS dataset at (e) pre-inoculation, (f) after inoculation, (g) flowering, and (h) senescence. Vertical bars represent intersection sizes (shared ASVs among treatment combinations), with the largest intersection highlighted in blue. Horizontal bars indicate the total number of ASVs within each treatment.

**S.F. 4.** Volcano plots showed log fold change against –log adjusted p-values. ASVs with adjusted p < 0.05 and log fold change > 1 or < −1 were classified as significantly up– or down-regulated, respectively. Volcano plots based on 16S rRNA amplicon sequencing data showing differential abundance analyses at the genus level comparing the control with each treatment across four sampling time points: pre-inoculation, after inoculation, flowering, and senescence. Each point represents a bacterial genus, plotted according to log2 fold change and statistical significance (−log10 P-value). Genera significantly enriched in the treatment relative to the control are shown as upregulated (red), whereas genera significantly depleted are shown as downregulated (blue). Genera not showing significant differential abundance are displayed in grey.

**S.F. 5.** Volcano plots showed log fold change against –log adjusted p-values. ASVs with adjusted p < 0.05 and log fold change > 1 or < −1 were classified as significantly up– or down-regulated, respectively. Volcano plots based on ITS rRNA amplicon sequencing data showing differential abundance analyses at the genus level comparing the control with each treatment across four sampling time points: pre-inoculation, after inoculation, flowering, and senescence. Each point represents a fungal genus, plotted according to log2 fold change and statistical significance (−log10 P-value). Genera significantly enriched in the treatment relative to the control are shown as upregulated (red), whereas genera significantly depleted are shown as downregulated (blue). Genera not showing significant differential abundance are displayed in grey.

**S.F. 6.** Bulk soil compared to rhizosphere of potato plants at the first sampling time point (1st of July). A-D, the bacteria rhizosphere microbiome. E-H, the fungal rhizosphere microbiome

**S.F. 7.** Bulk soil compared to rhizosphere of potato plants at the senescent time point (2nd of September). A-D, the bacteria rhizosphere microbiome. E-H, the fungal rhizosphere microbiome

**S.F. 8.** Differential abundance analysis was performed using the DESeq2 package [82] in R to identify microbial ASVs significantly affected by biocontrol treatments relative to the control. DESeq2 normalization was performed using the median-of-ratios method, and ASV counts were modeled using a negative binomial generalized linear model to account for overdispersion in sequencing data.

**S.T. 1**. Summary of co-occurrence network parameters derived from 16S rRNA gene data, including the number of nodes and edges, edge-to-node ratio, number and proportion of positive and negative edges, average clustering coefficient, and modularity.

## Notes

https://doi.org/10.5281/zenodo.23471271

### Competing Interest Statement

The authors have declared no competing interest.

