## Supplementary figures and images for "Building Bridges Belowground: *Pythium oligandrum* En-hances Rhizosphere Network Connectivity"

### Supplementary figure 1

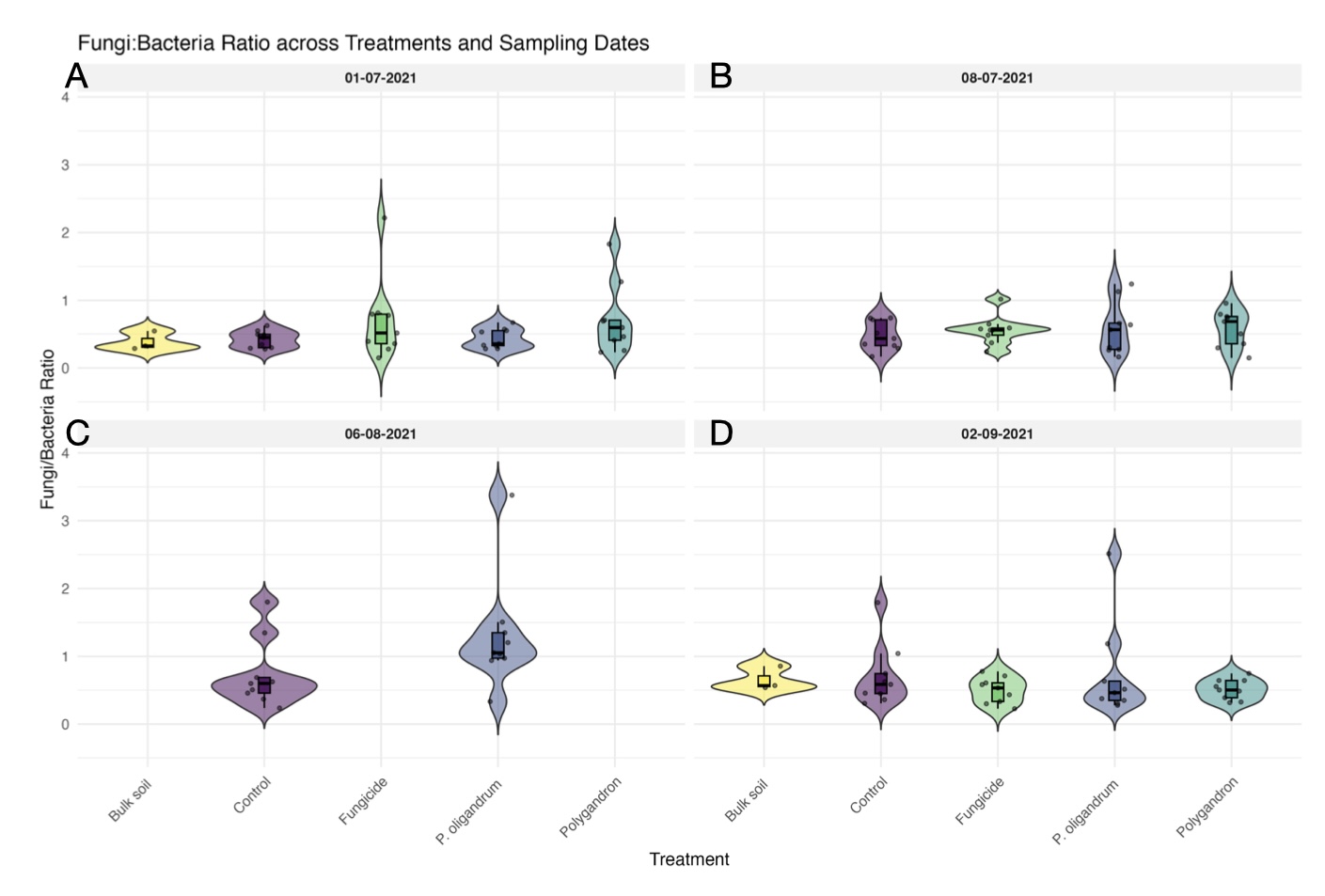

### Supplementary figure 2

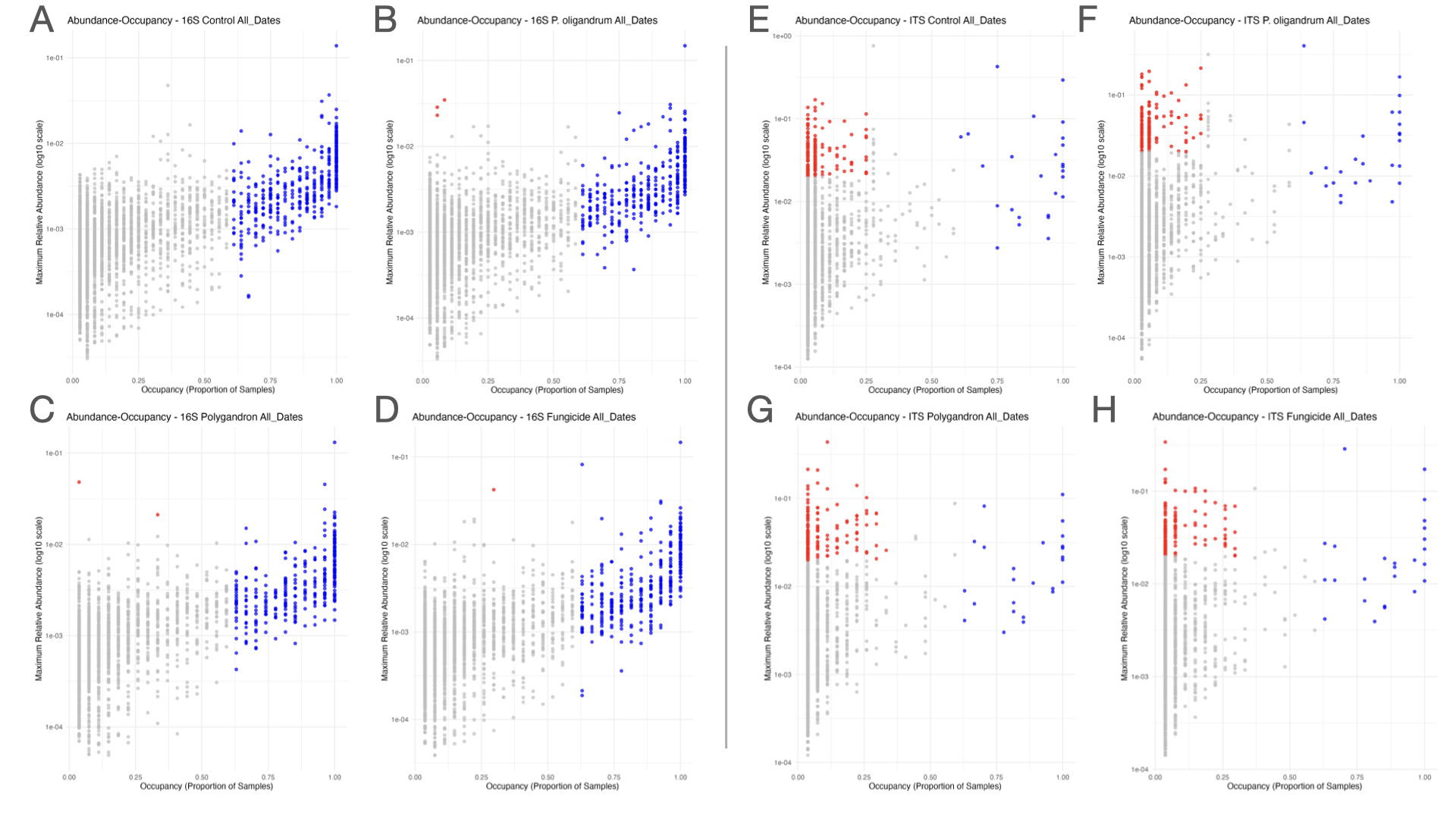

### Supplementary figure 3

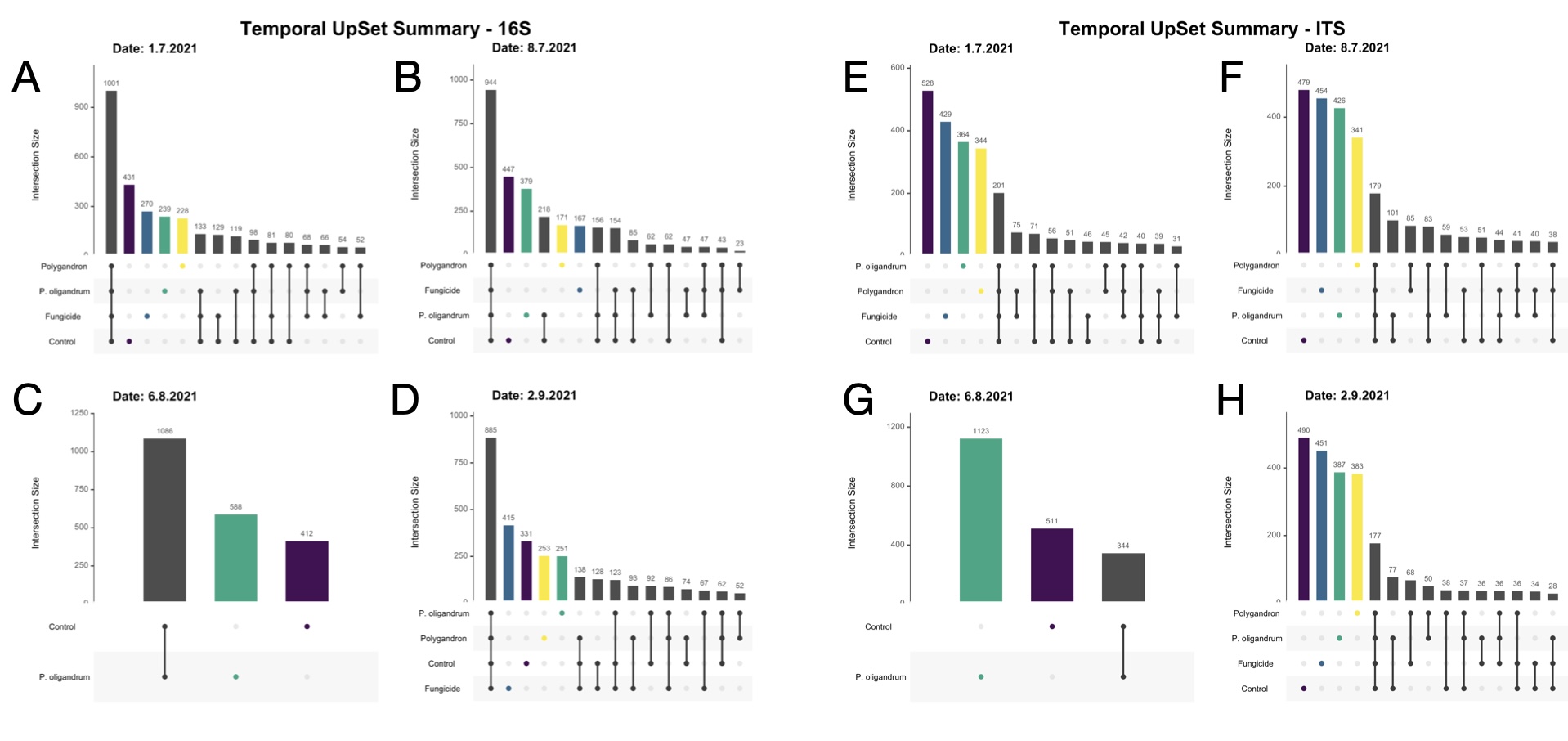

### Supplementary figure 4

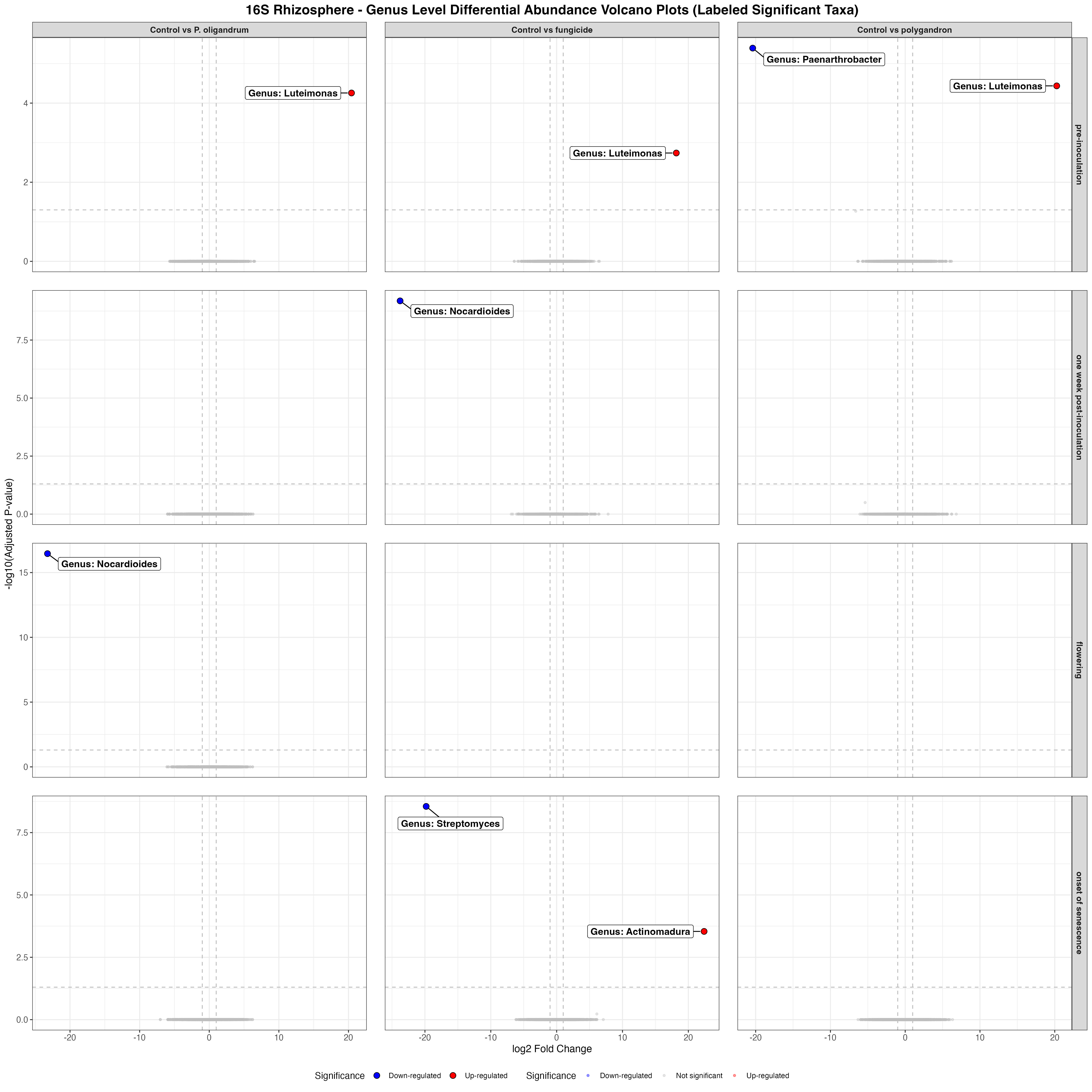

### Supplementary figure 5

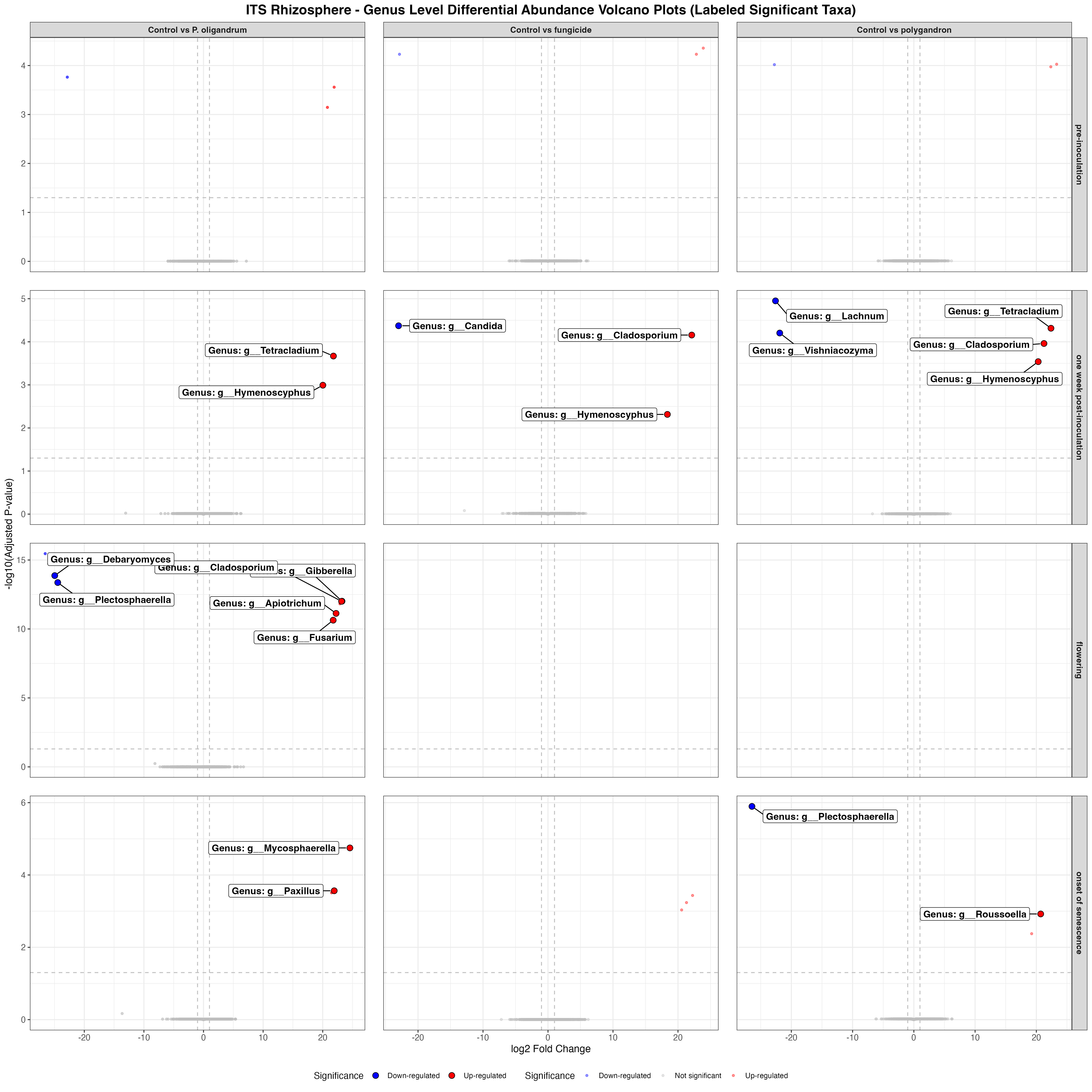

### Supplementary figure 6

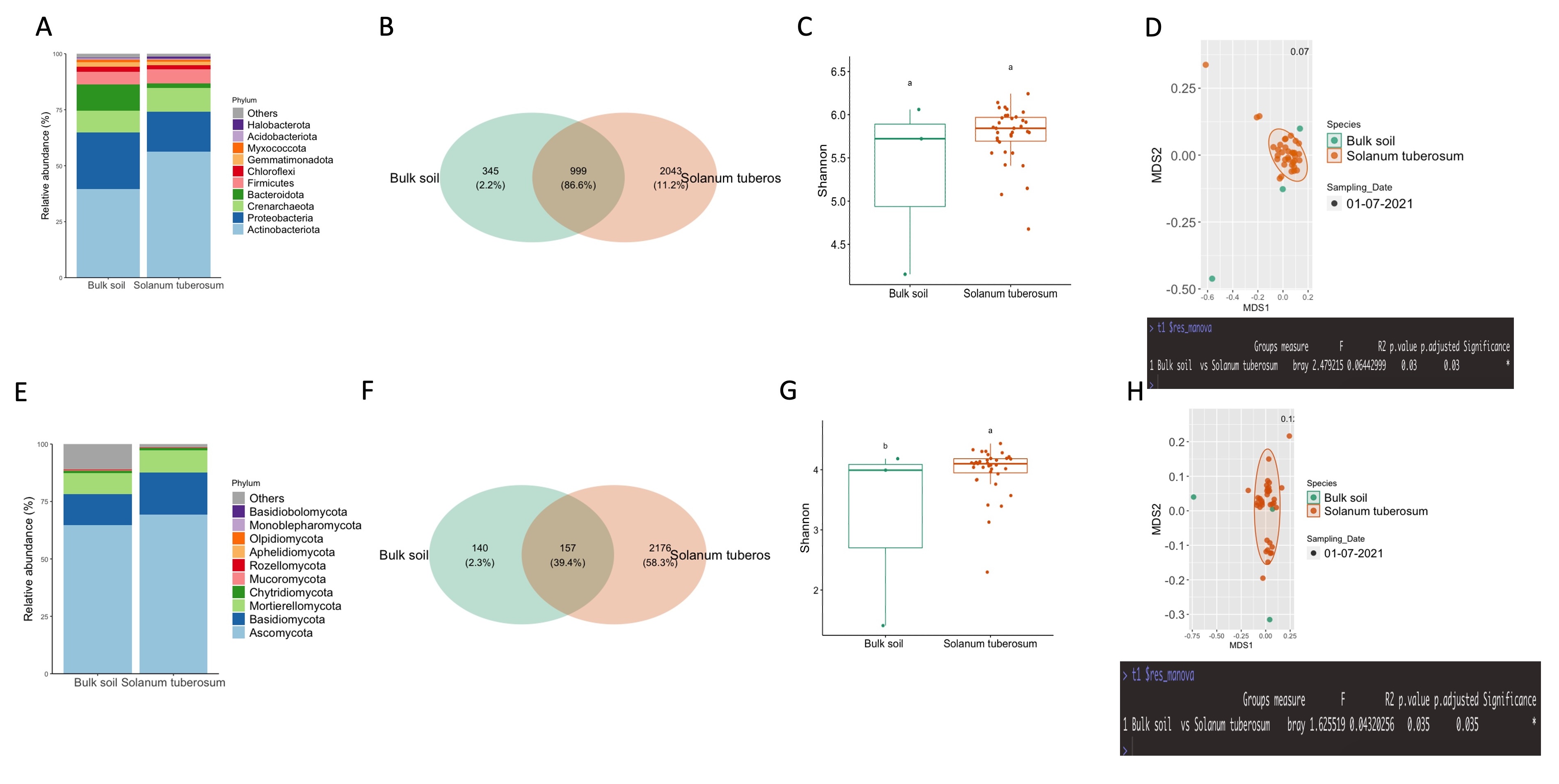

### Supplementary figure 7

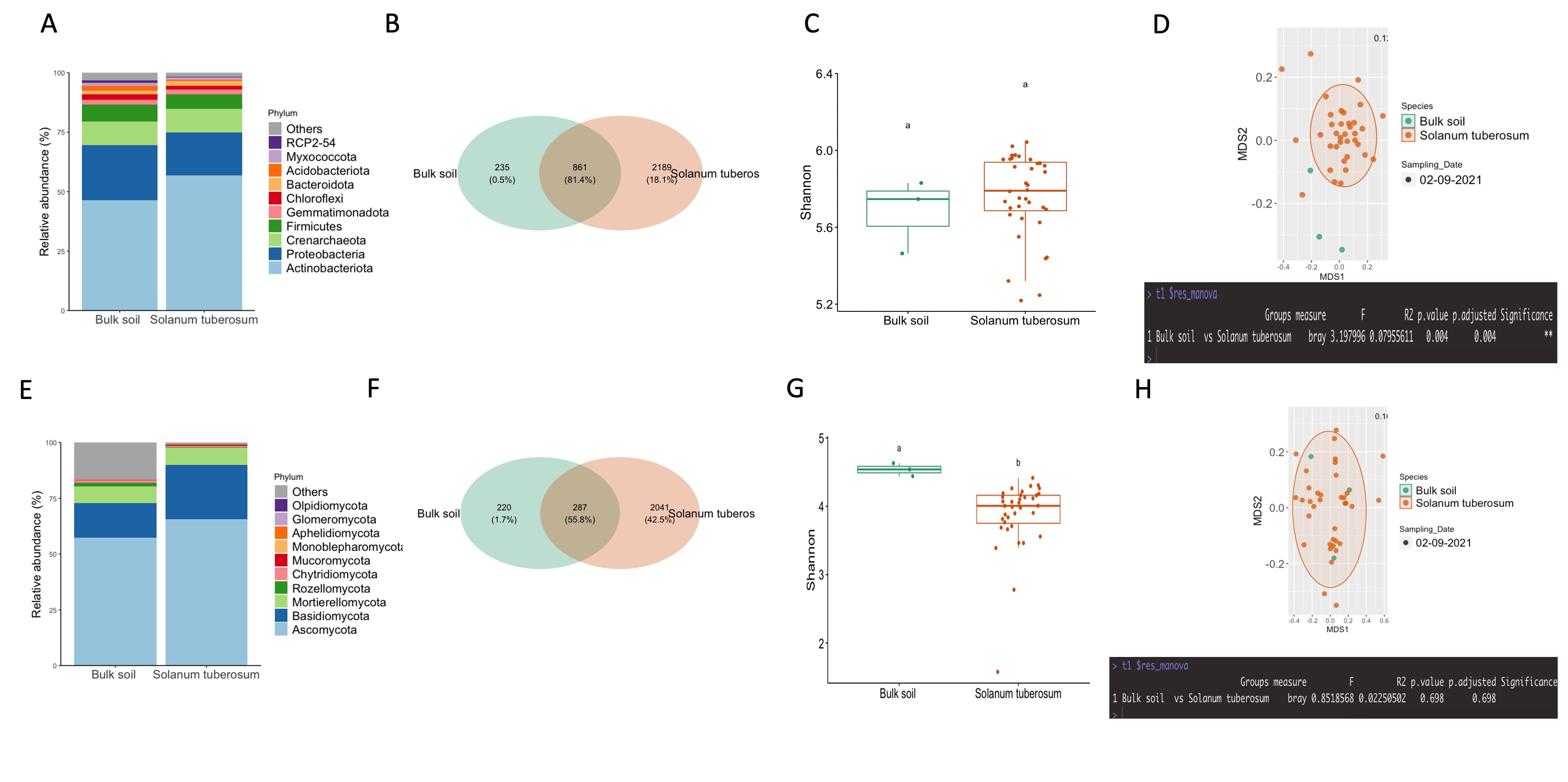

### Supplementary figure 8

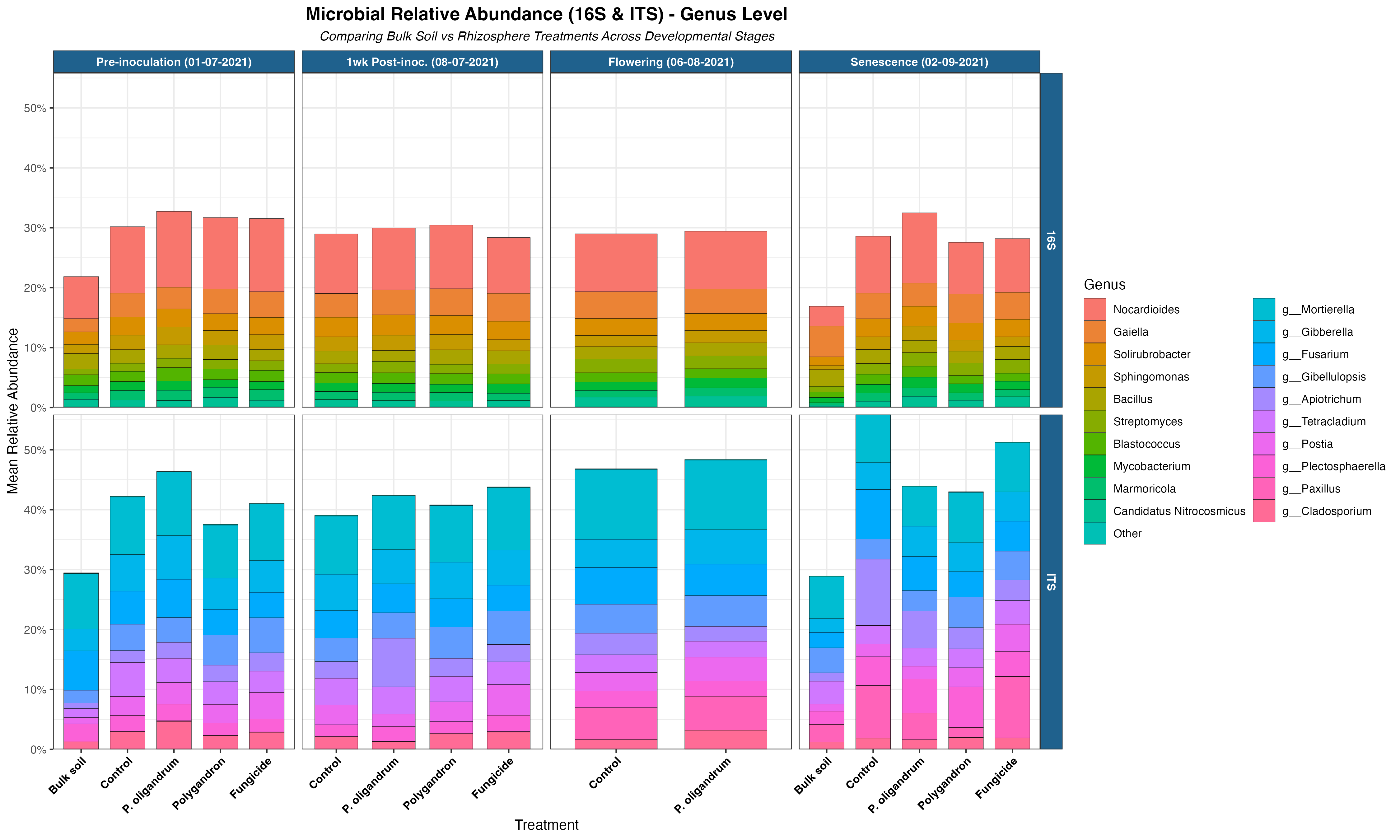

### Supplementary figure 8

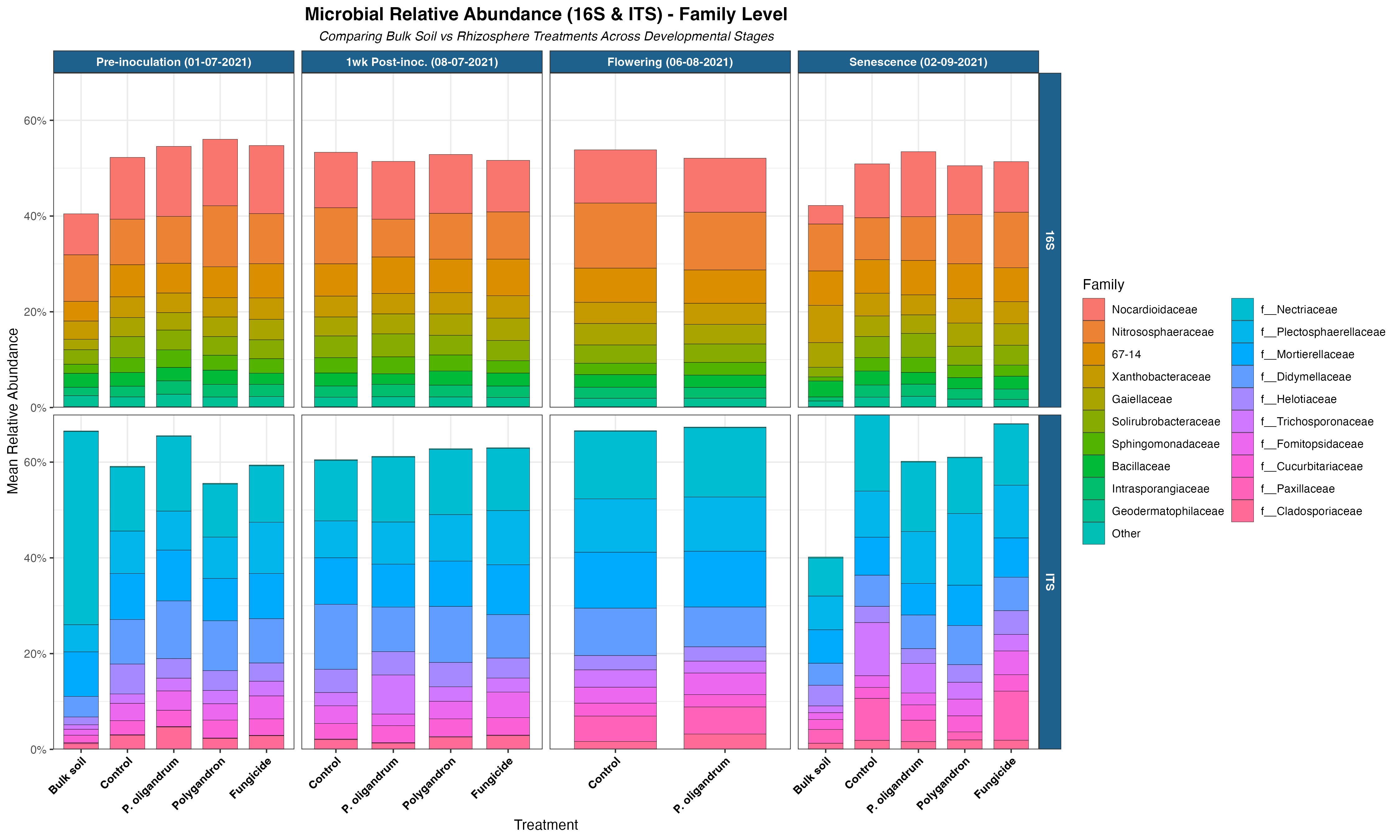

### Supplementary figure 8

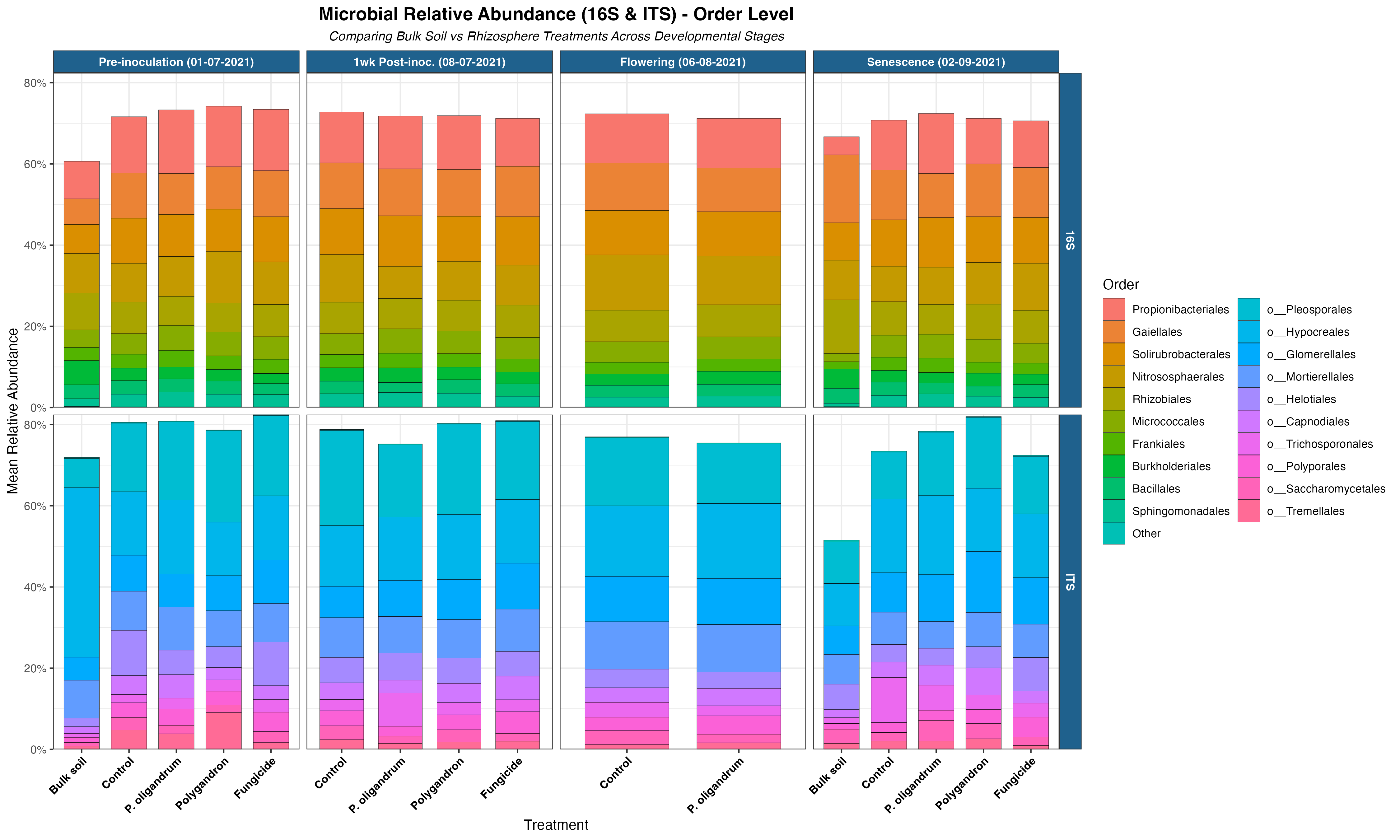

### Supplementary figure 8

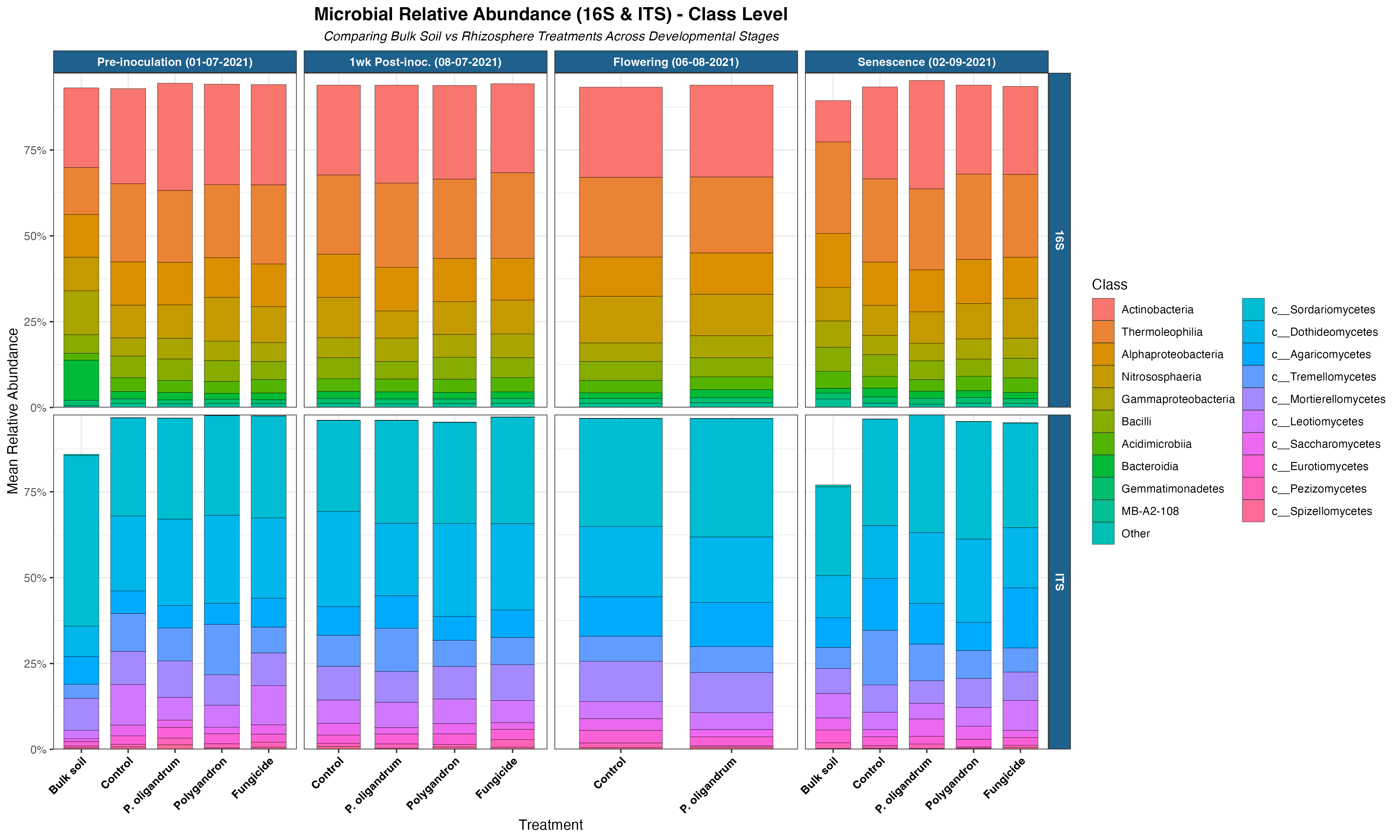

### Supplementary figure 8

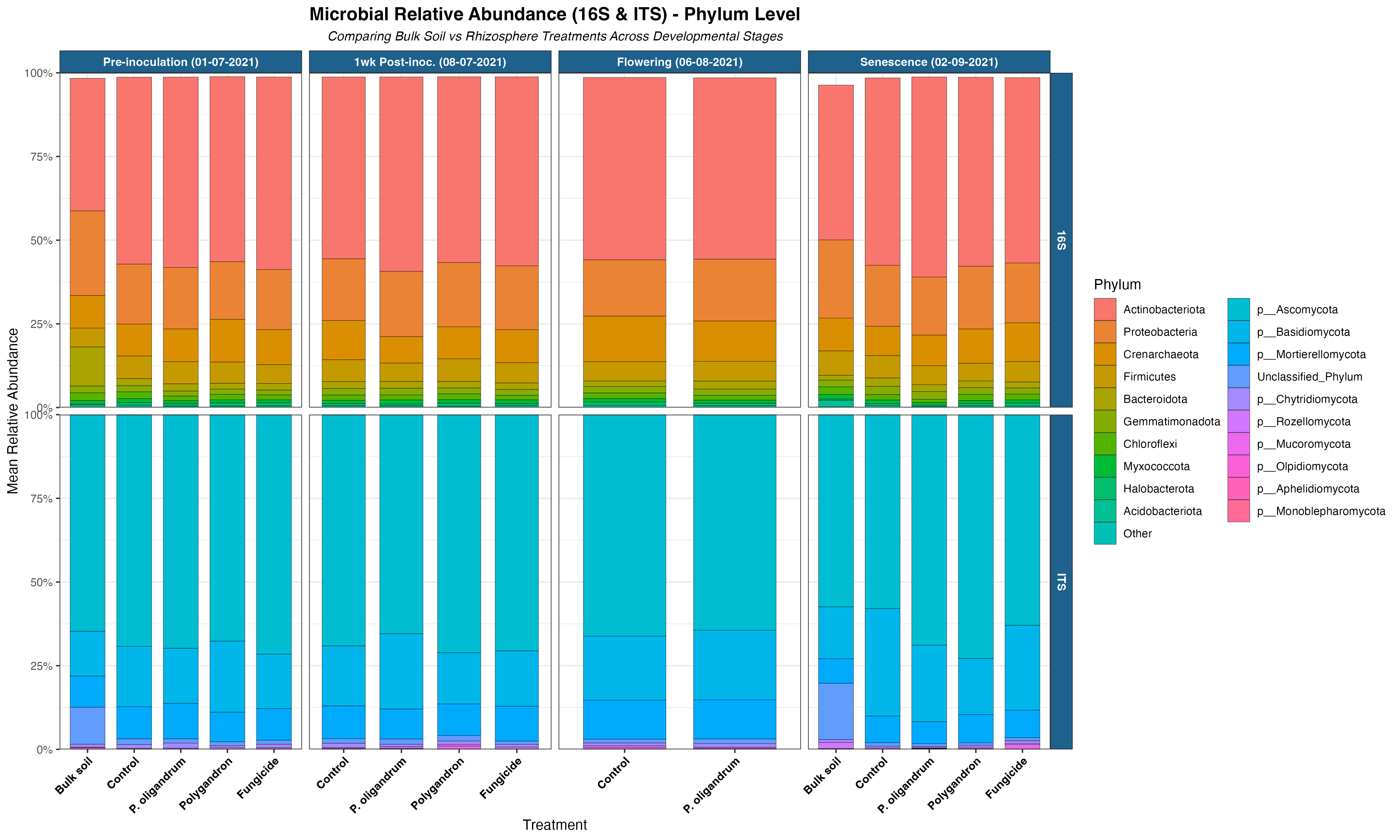

### Visual abstract

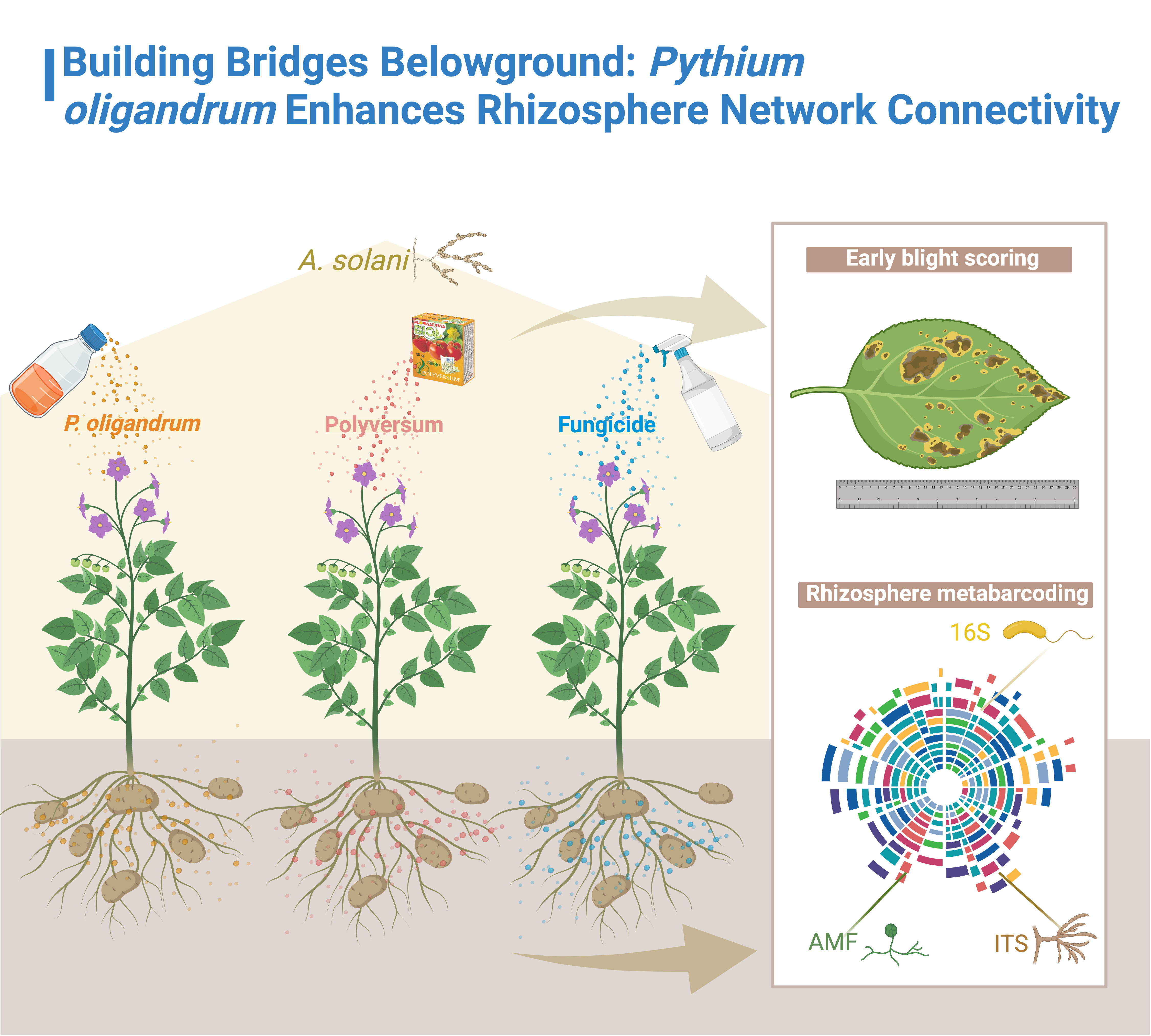
